# Mzion enables deep and precise identification of peptides in data-dependent acquisition proteomics

**DOI:** 10.1101/2023.01.17.524387

**Authors:** Qiang Zhang

## Abstract

Sensitive and reliable identification of proteins and peptides pertains the basis of proteomics. We introduce Mzion, a new database search tool for data-dependent acquisition (DDA) proteomics. Our tool utilizes an intensity tally strategy and achieves generally a higher performance in terms of depth and precision across twenty datasets, ranging from large-scale to single-cell proteomics. Compared to several other search engines, Mzion matches on average 20% more peptide spectra at tryptic enzymatic specificity and 80% more at no enzymatic specificity from six large-scale, global datasets. Mzion also identifies more phosphopeptide spectra that can be explained by fewer proteins, demonstrated by six large-scale, local datasets corresponding to the global data. Our findings highlight the potential of Mzion for improving proteomic analysis and advancing our understanding of protein biology.

## Introduction

Confident characterization of peptides and proteins from mass spectrometry (MS) data is essential for biological researches that employ MS-based proteomics techniques. Among a multitude of data interpretation facilities, closed searches of tandem mass (MS/MS) spectra from data-dependent acquisition (DDA)^1^ remains the baseline module in proteomics pipelines.^2–11^ Here, closed searches refer to experimental-to-theoretical matches with a predefined space in the variable modifications of theoretical peptide sequences.

Closed searches of DDA data compare MS/MS spectra to peptide sequences in protein databases.^12^ While search algorithms against the DDA data in general work well, qualitative and quantitative findings from software tools that match experimental to theoretical spectra do not always reconcile.^13^ Some of the discrepancy can be attributed to the different handling of users’ specified fixed and variable modifications by search engines. Others may be due to the scoring algorithms^14–17^ that arbitrate peptide spectrum matches (PSM) into the two domains of positive or false identifications, the boosting of machine data with complementary ions^18^, the size of search space, unexpected side effects of software, etc.

In this work, we present a tool, Mzion, that supports DDA searches of MS/MS spectra (Fig. 1). The tool matches separately the main sequences of MS/MS ions (e.g., *b*- and *y*-ions) and the satellite series (*b*^0^, *y*^0^, *b*^*^, *y*^*^ and doubly charged *b*^2^, *y*^2^, *b*^20^, *y*^20^, *b*^2*^, *y*^2*^). It then refrains the information learned from the satellite matches by disallowing them from being counted as independent evidences in peptide scoring. Instead, the satellite intensities are tallied onto the corresponding main fragment-ion intensities, followed by intensity-based enrichment analyses. Mzion enables generally deeper proteome discovery at higher or comparable precision, demonstrated by twelve large datasets. It also generalizes the handling of fixed and variable modifications that are sometime incompatible, with an additional benefit in search space minimization.

## Results

### Fixed and variable modifications

Interactions of fixed and variable modification terms specified in a database search may lead to additive effects that violate the fundamentals in chemistry. For instance, the accumulated modification from the interaction of fixed peptide N-terminal TMT10plex labeling and variable protein N-terminal acetylation may be undesirable. To address this, Mzion begins by deploying a minimal set of compatible fixed and variable modifications from a users’ specification (Supplementary Note 1). In the example of a users’ specification of fixed Carbamidomethyl (C), TMT10plex (N-term) and TMT10plex (K) and variable Acetyl (Protein N-term), Gln->pyro-Glu (N-term = Q), Oxidation (M) and Deamidated (N) (Supplementary Table 1), the compilation and minimization by Mzion leads to a total of twelve combinations of fixed and variable modifications (Supplementary Table 2).

It has been observed that an enlarged search space can reduce the numbers of identified PSMs and peptides when controlling false discovery rates (FDR).^19^ The combinatorial minimization by Mzion guards against undue expansion of a search space with regard to users’ intention in finding peptides that are more likely to present in samples. In an example of acetylome (Supplementary Table 3: dataset Lung_A), the numbers of PSMs, peptides (without modifications) and sequences (with applicable modifications) increase by 6.2%, 5.2% and 7.3%, respectively, upon the refinement of search space (Supplementary Figure 1). Analogously in an example of ubiquitylome (dataset Lung_U), the numbers of PSMs, peptides, sequences increase by 3.2%, 4.3% and 5.3%, respectively.

### Intensity tally from two-stage searches

A strategy of two-stage search is employed by Mzion. After matching the main sequences of MS/MS ions (e.g., *b*- and *y*-ions), the search engine then matches the satellite series of fragment ions (*b*^0^, *y*^0^, *b*^*^, *y*^*^ and doubly charged *b*^2^, *y*^2^, *b*^20^, *y*^20^, *b*^2*^, *y*^2*^), tallies their intensities to the corresponding main fragment ions, followed by intensity-dependent enrichment analyses of probability scores against the main fragment ions (Fig. 2a and **Methods**). In an example of phosphoproteome (dataset JHU_P2), the strategy led to a 6.5% increase in sequence identifications (Supplementary Figure 2).

The findings of PSMs are stored in files psmC.txtand psmQ.txtwhere the former contains the complete list of PSMs in the search space at a given set of search arguments and the latter gives the quality subset that passes a user-specified FDR threshold.

### High performance in global peptide identification with Mzion

We first compared the performance of Mzion to several other search engines using six datasets of 10-plex global TMT (dataset WHIM_G). Under the assumption of tryptic enzymatic specificity, all engines perform comparably with similar numbers of significant PSMs, peptides, sequences and proteins (Fig. 2b). Yet notable difference remains, exemplified by a relatively low percentage of common PSMs across search engines (Fig. 2c, Supplementary Notes 2 and 3). Within the six datasets, Mzion is in general the top performer in term of PSM, peptide and sequence identifications. It also regularly yielded the fewest numbers of proteins with single-peptide identifications (Supplementary Fig. 3). Analogous traits can be established with a pancreatic cancer dataset Panc_G1 (Supplementary Fig. 4).

We next tested Pearson correlations between the first two WHIM2 replicates in each of the six global datasets. Note that data stretching by extreme values can amplify correlations (Supplementary Note 4). We quantified additionally the data similarity by measuring the Manhattan distance of log2FC (less affected by reference choices) between replicated samples. We observed that Mzion confers high correlations and short similarity distances that are comparable to other search engines (Fig. 2d, Supplementary Fig. 5a-b).

We further assessed the precision in peptide and protein identifications against the global datasets using an entrapment strategy.^20^ Different to the estimates of FDR using a reversed database,^21^ the entrapment approach introduced an intermediate step by appending a low homology database to a target database. The combined database was then reversed and served as a decoy for FDR controls. Under 1% FDR of proteins, we observed that the entrapment rates of Mzion progressed comparably to the other four search engines (Supplementary Note 5). We also applied an orthogonal entrapment strategy by specifying the variable modification of TMTzero™ (TMT0) to peptide N-terminal and site K in the search. The molecular mass of TMT0 differs to that of TMT10plex by 5.01045 Da and are expected to be absent in general from the WHIM_G dataset. Among the five search engines, Mzion yielded low entrapment rates that progress consistently along the decreasing peptide scores (Supplementary Fig. 5c).

Identities of semi/non-tryptic peptides are among the so-termed dark matter in MS data.^22^ We further searched the WHIM_G dataset at no enzymatic specificity (NES). We found that, in general, Mzion outperforms other search engines by numbers of PSMs and peptides at higher or comparable precision (Supplementary Fig. 6-7, Supplementary Table. 4). For instance with the JHU_G2 subset, Mzion identified twice as many PSMs, peptides and sequences than the search engine that reported the second highest correlations (Fig. 2b). Presumably the production of semi/non-tryptic peptides are often less defined than tryptic ones. In line with the notion, decreased correlations and increased distances between replicated samples were observed for all search engines when comparing the semi/non-tryptic to the tryptic findings (Fig. 2e).

### Deeper proteome coverage in phosphopeptides

Protein phosphorylation is one of the most widely explored post-translational modifications in proteomics. We demonstrate the performance of Mzion in the characterization and quantitation of immobilized affinity chromatography (IMAC) enriched phosphopeptides, using the six datasets of 10-plex TMT under WHIM_P. We noted that Mzion in most cases is the leading search engine in PSM, peptide and sequence identifications (Supplementary Fig. 8a). We further observed that Mzion yielded short similarity distance between replicated samples, when compared to the search engine that reported the second highest depth in phosphopeptides (Supplementary Fig. 8b). In the example of the JHU_P2 dataset, Mzion characterized 10.1% more peptides than the search engine that yielded comparable entrapment rate, and yet at higher correlation and shorter similarity distance (Fig. 3). Interestingly, the greater number of peptides can be explained by 11.1% fewer proteins.

It is worthwhile to note the greater discrepancy in the identities of sequences than peptides. For instance, 26.6% of sequences versus 7.3% of peptides were missed from Mzion when compared to the findings from Mascot (Figure 3). The observation is consistent with the documentation of low correlations in the phosphorylation site specificity between search engines.^23^ The localization probability by Mzion is modified from the AScore algorithm.^24^ Instead of assuming binomial probabilities, Mzion applied alternatively a counting statistics to weigh the localization probability.^25^ For instance, at a 2:1 counts of localization-specific observations, the former will be given a 0.67% probability in localization. Analogous at a 1:0 ratio, the former will be 100% responsible for the site specificity. Different to the binomial model in AScore, the localization score with Mzion will be the same at either ratio 1:0 or 2:0.

We further tested the accuracy of phosphopeptide identification with Mzion using dataset Ferries_spikes. The dataset comprises 191 spiked phosphopeptide sequences with variations in phosphosites. We found that all search engines performed comparably in the numbers of identified phosphopeptides and sequences (Supplementary Table 5). Among the five engines, Mzion identified 177 phosphosequences, a number that is fewer than the expected value (of 191).

### General purpose searches with Mzion

Mzion is a general purpose search engine currently for closed searches of tandem mass spectra. For instance, it is readily applicable to feature matching of acetylome and ubiquitylome, as well as SILAC experiments (Supplementary Table. 6). In the examples of small datasets from single-cell proteomics, we observed that all search engines performed comparably with Mzion reported the fewest proteins at single-peptide identifications (Supplementary Fig. 9).

## Discussion

We have shown that Mzion is capable of in-depth characterization and quantitation of peptides and proteins from DDA data. We further showed that Mzion regularly captures DDA peptides and proteins at high precision. The depth and precision may facilitate biological research by generating novel insights with reliable proteomic discovery.

The outputs of Mzion (psmC.txt) contains both target and decoy findings and can be readily coupled to post-search algorithms, such as Percolator for re-scoring.^26^ In the example of Percolator post-processing of dataset JHU_G2, 38,804 PSMs that can be assigned to 27,988 peptides and 31,691 sequences were characterized additionally using the two predictors of precursor mass errors and retention times. To avoid the probable confusion that the gain by Mzion algorithm is due to post processing, we restrict ourselves in this work to assess the kernel performance of Mzion without Percolator. The quality output of Mzion, psmQ.txt, is derived from the complete output, psmC.txt. Comprehensive analyses of Mzion outputs are, however, performed with proteoQ(http://github.com/qzhang503/proteoQ) with additional measures in quality metrics and capabilities in bioinformatics.

Part of the reason for the popularity of DDA may be ascribed to its compatibility to high-throughput, multiplex TMT workflows. Data independent acquisition (DIA) methods, on the other hand, allow all precursors in a wide isolation window of *m/z* over a wide dynamic range of intensity being fragmented simultaneously for MS2 interrogation.^27^ It may seem intuitive that spectrum simplifications analogous to the intensity tally by Mzion may, to some extents, aid the separations of signals from background noises in more complex DIA spectra,^18,28–31^ and reduce the chance of matching everything to everything. However, the potential benefits are yet to be explored, especially with regard to DIA signals at low MS2 intensities within a given isolation window. A future direction with Mzion may include additional DIA scoring algorithms, aided by spectrum simplification.

Mzion is developed under the R software environment: an interpreted programming language that is commonly used in statistical computing and data analysis. The language may not yet be rated among the fastests for reasons such as limited supports of reference semantics. In spite, with the broad community efforts, many R libraries are rooted in compiled languages such as C and C++. The underlying mechanisms allow Mzion to execute suitably in proteomics database searches where the speed of analysis is an important dimension in tool performance. In the example of dataset BI_G1 (number of RAW files: 25; databases: Refseq human mouse and cRAP; enzyme: trypsin/P; number of missed cleavages: 4; precursor mass error tolerance: 20 ppm; product ion mass error tolerance: 20 ppm, fixed and variable modifications: Supplementary Table 2), the forward and reversed matches of *b*- and *y*-ions are typically done in about 65 mins (the complete process in about 101 mins or 97 mins with cached precursor masses) when using an 8-core (Intel i7-7820x CPU, 3.6 GHz) PC at 32 GB RAM. The corresponding search times are approximately 85 mins with Mascot (without post-processing), 500 mins with MaxQuant, 45 mins with MSFragger and 220 mins with MS-GF+. Currently the search speed of Mzion is limited by the generation of theoretical MS/MS ions that are subject to heavy permutations of variable modifications and sites (e.g. searches of phosphopeptides). We conceive further performance improvement upon rewriting the permutations in C++ or by adapting pre- computed permutations.

## Methods

### Workflow of Mzion

#### Software environment and input data

Mzion was developed under the free software environment R^32^ and Posit/RStudio^33^. It processes peak lists from MSConvert^34^ for searches of experimental mass spectra. When converting RAW MS files, the peak lists can be at either Mascot generic format (MGF) or mzML format. The option of TPP compatibility needs to be checked (as defaulted) at either format for downstream parsing of the title lines in the peak lists with proteoQ (http://github.com/qzhang503/proteoQ). With mzML outputs, the default of Use zlib compressionneeds to be *unchecked* (opposite to the default) for proper decoding of binary data to MS2 *m/z* and intensity values.

#### Compilation of fixed and variable modifications

The same site, including “N-term” or “C- term” of a peptide, at multiple variable modifications are allowed (for example variable TMT10plex modification and acetylation both at site K). However, the same site being specified under both fixed and variable modifications are considered initially incompatible in that the addition of the variable mass to the fixed mass may be chemically unsound (for example fixed TMT10plex modification and variable acetylation of K). To handle this, Mzion first coerces the incompatible fixed modifications, *F*_*X*_, to the category of variable modifications. Sets of combinatorial variable modifications are then computed. Among them, combinations containing multiple “N-term” or “C-term” modifications are removed (for example TMT10plex modification at “N-term” and acetylation at “Protein N-term”). The program next looks for a coerced site at position “Anywhere” and removes the combinations without the site (for example removals of combinations lacking K with the fixed-to-variable coercion of TMT10plex modification of K). The same rule applies to coerced terminal modifications. In case of multiple coercions of “Anywhere” sites, combinations that do not contain *all* of the coerced sites are removed. Finally for each combination, the coerced *F*_*X*_ are reverted back to fixed modifications if there are no site violations (for example *F*_*X*_ is TMT10plex modification at “N-term” and there is no other variable “N-term” modifications). Note that to obtain an additive effect, a new Unimod^35^ entry containing multiple modifications to the same site can be constructed (for example TMT10plex modification at “N-term” and the formation of pyro-glutamic acid from N-terminal glutamine). The Mzion utility add_unimodcan be used for the purpose.

#### Amino-acid look-ups

Look-ups of amino-acid residues are stored as named vectors for each combination of fixed and variable modifications (for example, twelve modules of look-ups when instantiating the modifications specified in Supplementary Table 2). Sites and variable modifications are in names and the corresponding mono-isotopic masses are in values. For example with the specification of variable “Oxidation (M)”, both unmodified “M” and “Oxidation (M)” are available in the look-up at masses of 131.0405 and 147.0354, respectively. Masses of fixed modifications are applied directly to the corresponding sites. Titles, positions and sites of modifications are linked to the look-ups as R-language attributes. Additional attributes include sites of coercion, sites with neutral losses, etc. The attributes were calculated once and can be re-used when needed. For instance when calculating the theoretical MS2 ions, an attribute of “nl+” prompts the calling of utilities that consider the permutation of neutral losses”.

#### Peptide scores

Experimental tandem spectra are first searched against theoreticals by primary ions. Additional searches are performed against tandem spectra with a minimum of *n* matched primary *b* and/or *y* ions (*n* = 6 by default). Experimental intensities of the matched secondary ions are ascribed to those of the corresponding primary ions, for example,

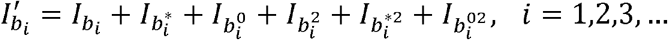

where *I* stands for experimental intensity and 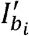 is the boosted experimental intensity upon summing 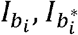 etc. Theoretical *m/z* are next ordered decreasingly according to the boosted intensity.

| $m/z$ (Expt) | Intensity (Expt) | $m/z$ (Theo) | $k$ | $x$ | $p$ |
| --- | --- | --- | --- | --- | --- |
| 248.1809 | 88,320 |  | 14 | 0 |  |
| 376.2761 | 81,536 | 376.2757 | 15 | 1 | 0.3460 |
| 863.4354 | 65,913 | 863.4348 | 16 | 2 | 0.1077 |
| 305.1691 | 65,251 |  | 17 | 2 |  |
| 551.8666 | 59,457 |  | 18 | 2 |  |
| 396.2363 | 54,537 |  | 19 | 2 |  |
| 291.1717 | 49,432 |  | 20 | 2 |  |
| 551.8262 | 49,064 |  | 21 | 2 |  |
| 331.2138 | 48,669 |  | 22 | 2 |  |
| 551.6346 | 46,109 |  | 23 | 2 |  |
| 505.3192 | 45,770 | 505.3183 | 24 | 3 | 0.0520 |
| 470.3096 | 42,045 |  | 25 | 3 |  |
| 551.9680 | 41,824 |  | 26 | 3 |  |
| 343.2545 | 40,270 | 343.2543 | 27 | 4 | 0.0150 |
| 490.3232 | 38,931 | 490.3227 | 28 | 5 | 0.0028 |
| 552.3222 | 36,185 |  | 29 | 5 |  |

Hypergeometric probabilities (*p*) are assessed progressively against the numbers of matched primary ions (*x*) versus the numbers of MS2 features (*k*). The best (minimum) probability, *P*, is applied and the corresponding entrenchment scores is -10 - log_lO_(*P*). By default, the FDR is estimated by a target-decoy approach against the subclass of search results that yields the maximum number of matches. The estimated FDR are then transferred to all results.^36^

#### Protein groups

Identities of proteins and peptides are arranged in a sparse logical matrix (R package Matrix) with column and row associations to proteins and peptides, respectively. A logical value of 1 in the matrix stands for the presence of a peptide under a protein. The matrix is next divided rowly by unique (r1) and shared (r2) peptides (Figure 4). The submatrix r2 is further separated columnly by proteins with (c1) or without (c2) peptide counts. The same column separation applies to the submatrix of unique peptides (r1). Note that values in the submatrix intersecting r2 and c2 are all logical zeroes and each of the c2 proteins forms their own group. Further note that unique peptides have no effect on the grouping of proteins by distance.

It is sufficient to perform protein-peptide grouping against the submatrix intersecting c1 and r2, followed by additional increments in cardinal numbers for proteins with single peptide identifications. Pairwise distance of proteins are first assessed (R package proxyC) by the logical condition !any(A & B) (not any “both A and B are 1”). It returns 0 if protein A and B share peptides and 1 if not. The dichotomy distance matrix is next clustered hierarchically with the agglomeration method of single link. The cluster is then cut at an arbitrary height greater than 0 (heights are 0 between proteins that share peptides). Finally, essential proteins are determined using a greedy set cover algorithm.^37^

### Database searches

MS data were converted to peak lists using MSConvert (v3.0.22279). The MS/MS spectra were analyzed using Mascot (v2.8.0.1), MaxQuant (v2.0.2.0), MSFragger (v18.0), MS-GF+ (v20230112) and Mzion (v1.2.3). The top-100 most abundance product ions were used with MSFragger and Mzion. The allowance in mass error was set to +/-20 ppm for both precursor and product ions. A maximum of 4 missed cleavages was allowed for searches assuming the digestion enzyme of trypsin/P. The range of peptide lengths was between 7 and 40 residues and the range of precursor masses between 700 and 4500 Da (Supplementary Fig. 11). In multiplex TMT studies, the tolerance in reporter-ion mass error was set to +/-10 ppm with Mascot, MS-GF+ and Mzion, 0.003 Da with MaxQuant and +/-20 ppm with MSFragger. A maximum of five variably modified sites per peptide sequence was allowed with a maximum of three positions at the same modification and 64 permutations in positions.

#### Datasets WHIM_G and WHIM_P

The global WHIM_G dataset was set up to search against a Refseq database of human mouse proteins (ver Jul. 2018; 56,789 entries) and common contaminant proteins (cRAP, v1.0 Jan. 2012; 116 entries). The Refseq database was changed to Uniprot human mouse (ver Oct. 2020; 37,522 entries) when searching against the phosphopeptide WHIM_P dataset. Cysteine carbamidomethylation was specified as a fixed modification. TMT10plex modification of peptide N-terminals and K were specified under the quantitation method with Mascot and MaxQuant. The TMT10plex modification of K and peptide N-terminals were set as fixed modifications with MSFragger, MS-GF+ and Mzion. Methionine oxidation, asparagine deamidation, N-terminal glutamine to glutamic acid conversion and protein N-terminal acetylation were specified as variable modifications. Phosphorylation of S, T and Y were specified additionally as variable modifications when searching against the WHIM_P dataset. The specifications of variable asparagine deamidation and N-terminal glutamine to glutamic acid conversion were removed in the searches of WHIM_G at no enzymatic specificity. Results were filtered at 1% FDR at proteins (also met at the levels of PSM and peptide) with the exception of Mascot and MS-GF+ results being reported at 1% engine-specified PSM FDR. MSFragger PSMs were further processed with PeptideProphet.

Additional datasets. The Uniprot database was subset by species human (20,512 entries) when searching against the datasets of lung (acetylome Lung_A1_A10 and ubiquitylome Lung_U) and pancreatic (Panc_G1) carcinomas, as well as datasets Ferris_spike, single-cell Hela at 200pg and 2 ng inputs. Lysine acetylation (K-Ac) and carbamylation were both set to variable modifications in acetylome searches. Lysine ubiquitylation (K-GG) with and without TMT10plex were set to variable modifications in ubiquitylome searches. The search parameters against the WHIM_P were used for the searches of dataset Ferries_spike with the exclusion of TMT10plex modifications. The Dong_SILAC dataset was searched against a Uniprot database of E-coli (Jul. 2022; 4,564 entries) and the cRAP. The same fixed and variable modifications specified in the global WHIM_G searches were used except for TMT10plex.

Entrapment analysis by species was performed against the combined database of Refseq human mouse (or Uniprot human mouse with dataset WHIM_P), Uniprot arabidopsis thaliana (Sept. 2022; 16,312 entries) and cRAP with the identical sets of fixed and variable modifications specified in the WHIM_G (or WHIM_P) searches. Entrapment analysis of global data by TMT0 modifications was performed with the same settings in the WHIM_G searches, with the additional variable TMT0 modification to both K and peptide N-terminal.

#### Data analysis

Results from Mascot and MaxQuant were filtered by a minimum of 6 matching product ions (the same applied to MSFragger and Mzion via search arguments). Note that the tolerances in precursor and product-ion mass errors can be interpreted differently by search engines. Results of Mascot, MSFragger, MS-GF+ and Mzion were further filtered by +/- 10 ppm in precursor mass errors. With quantitative multiplex TMT data, PSMs without peptide N-terminal modifications were also removed from MSFragger (filtered by “N-term” under column “Assigned Modifications” in the psm.tsv output) and Mzion (automated with the latter when compiling fixed and variable modification sets). PSMs with all missing precursor intensity across samples were removed from quantitative analysis. Protein identifications were summarized to gene products.

### User-interface utilities

The main utility of Mzion is matchMS for database searches. Additional user-interface utilities include table_unimodthat tabulates titles, sites, positions and monoisotopic masses of all Unimod modifications, find_unimodthat extracts the information of a specific Unimod entry with additional specifications in neutral losses, parse_unimodthat checks the grammar of a user-specified modification, load_fasta2that loads FATSTA databases, calc_unimod_compmassthat calculates the monoisotopic mass of a modification by a chemical composition, add_unimod, remove_unimodand remove_unimod_titlethat add or remove Unimod entries. Utility mapMS2ionsmaps MS2 ions between theoreticals and experimentals and make_mztabprepares mzTab files for publication compliance. Utilities calc_monopeptideand calc_ms2ionseriescalculate the possible precursor masses and MS2 product ion series for a peptide sequence at a given set of fixed and variable modifications.

## Acknowledgements

QZ are grateful for the insightful comments from the anonymous reviewers. QZ thanks Drs. Hao Chi, Dirk Eddelbuettel, Lukas Käll, Matthew Monroe, Alexey Nesvizhskii, Clay Semenkovich, Reid Townsend and Kohei Watanabe for discussion.

## Data availability

The codes for Mzion is freely available at http://github.com/qzhang503/mzion under MIT license. All data generated or analysed during this study are included in the published articles summarized in Supplementary Table 3.

