## Supplementary Figures for "Mzion enables deep and precise identification of peptides in data-dependent acquisition proteomics"

**Supplementary Figure 1.** Mzion findings against datasets Lung_A (Acetyl) and Lung_U (Digly) at fixed or variable TMT10plex modifications (both K and peptide N-terminal). (F) fixed TMT10plex at K and peptide N-terminal with applicable coercions to variable modifications; (V) variable TMT10plex at K and peptide N-terminal. The first ten (A1 to A10) subsets were used for the acetylome analysis. All twenty subsets (U1 to U20) were used for the ubiquitylome analysis. The acetylome includes the modification of Carbamyl (K). The ubiquitylome includes the additive modification of TMT10plex+Digly (K). Results from fixed TMT10plex were subset to the space of modification groups in variable TMT10plex.

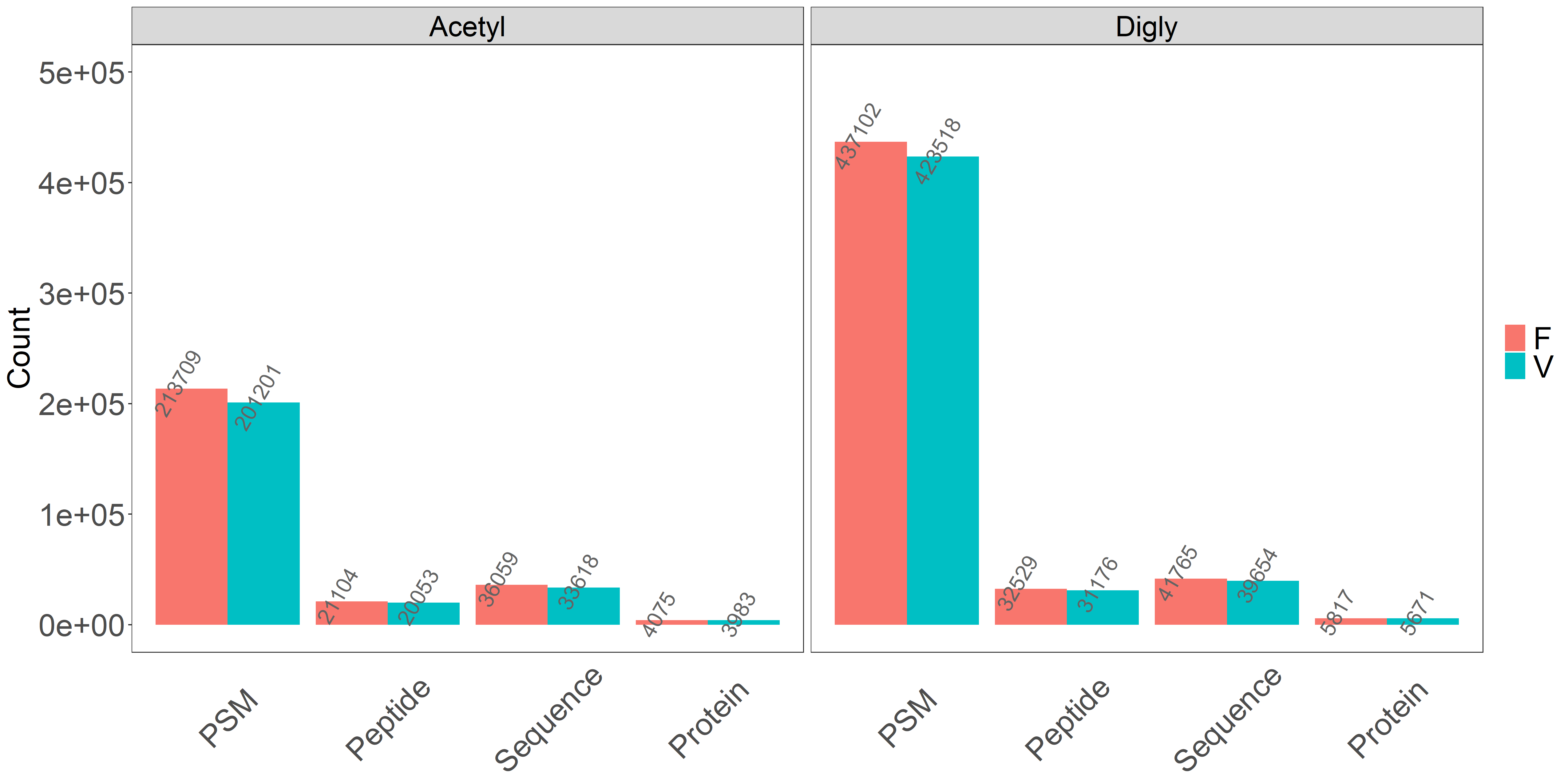

**Supplementary Figure 2.** Mzion findings with or without intensity tally from datasets (a) WHIM_G and (b) WHIM_P.

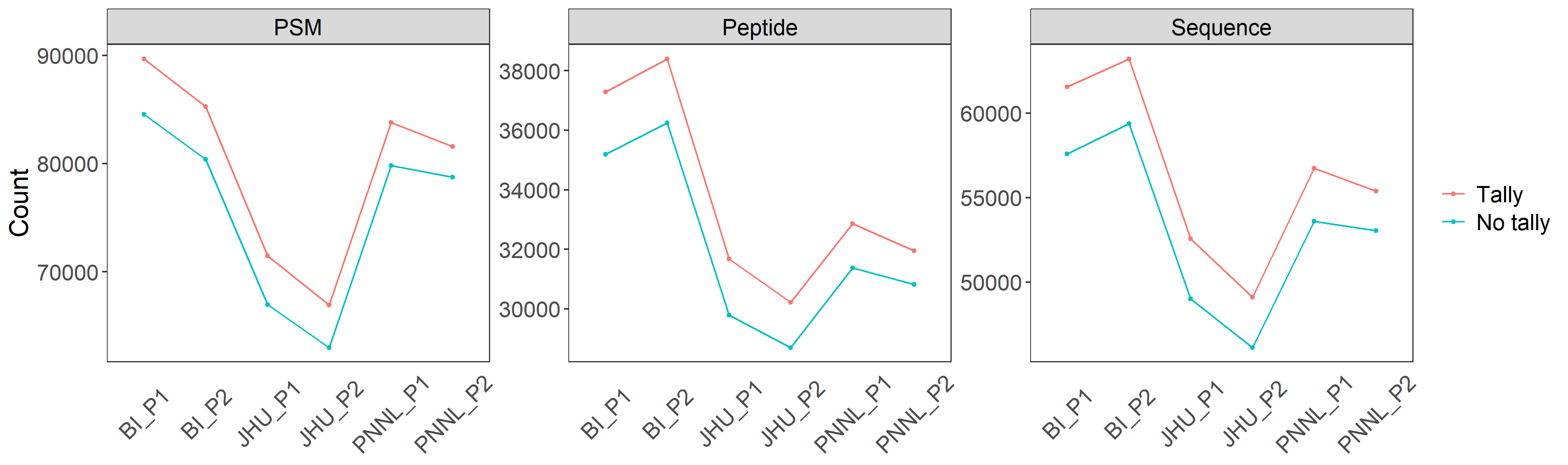

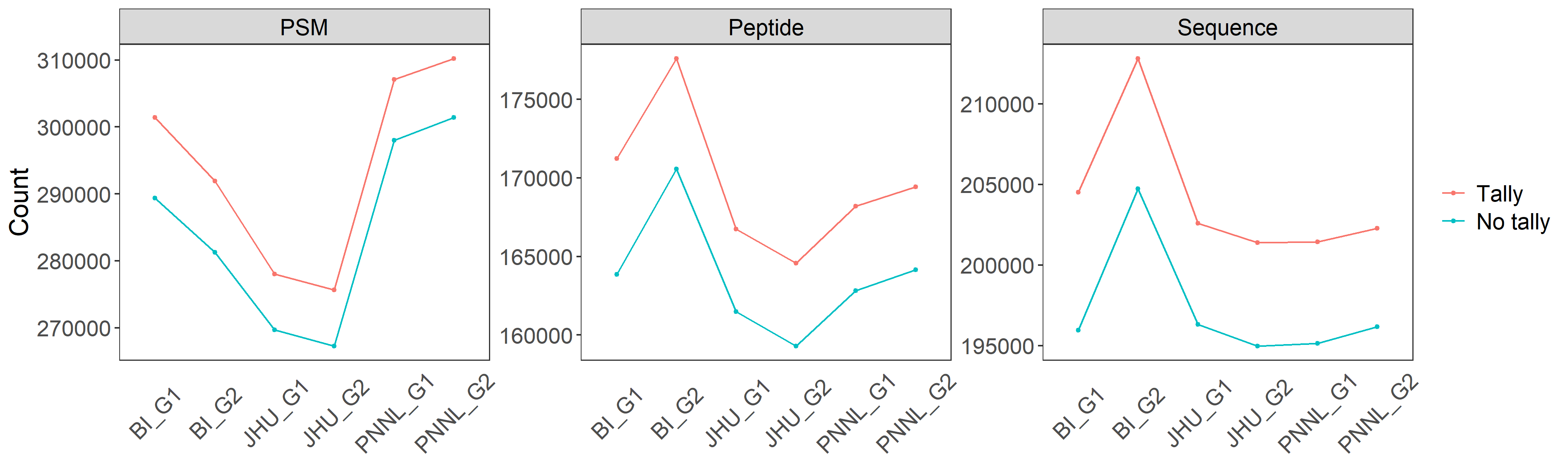

**b**

**a**

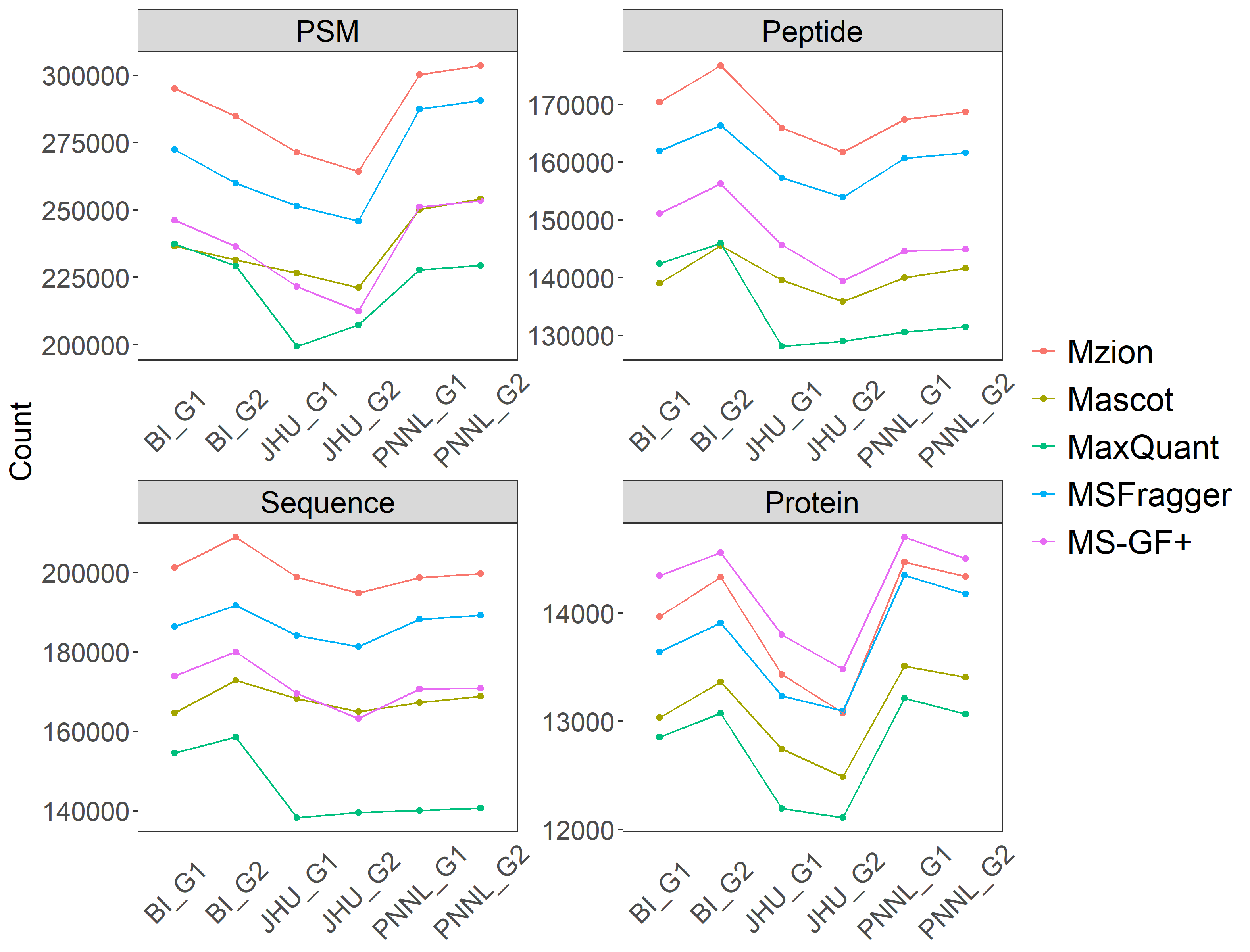
**Supplementary Figure 3.** Related to Figure 2. Tryptic findings from dataset WHIM_G. (a) Counts of PSMs, peptides, sequences and proteins. (b) Numbers of unique identifying peptides under proteins. Counts of proteins with one-peptide identification are shown in the insets.

**a**

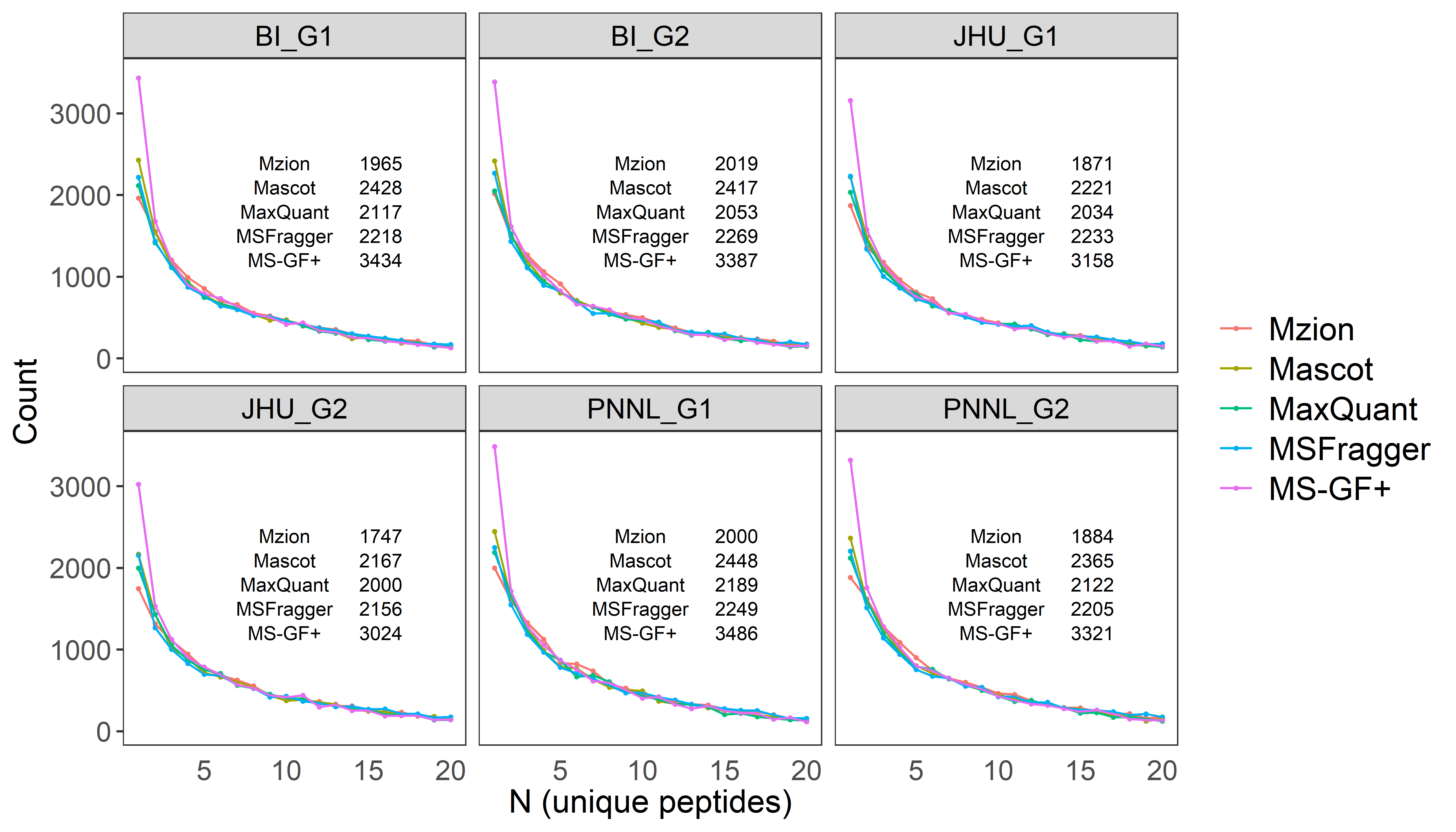

**b**

**Supplementary Figure 4.** Related to Figure 2. Tryptic findings from dataset Panc_G1. (**a**) Counts of PSMs, peptides, sequences and proteins. (**b**) Numbers of unique identifying peptides under proteins. Counts of proteins with one-peptide identification are shown in the inset.

**a**

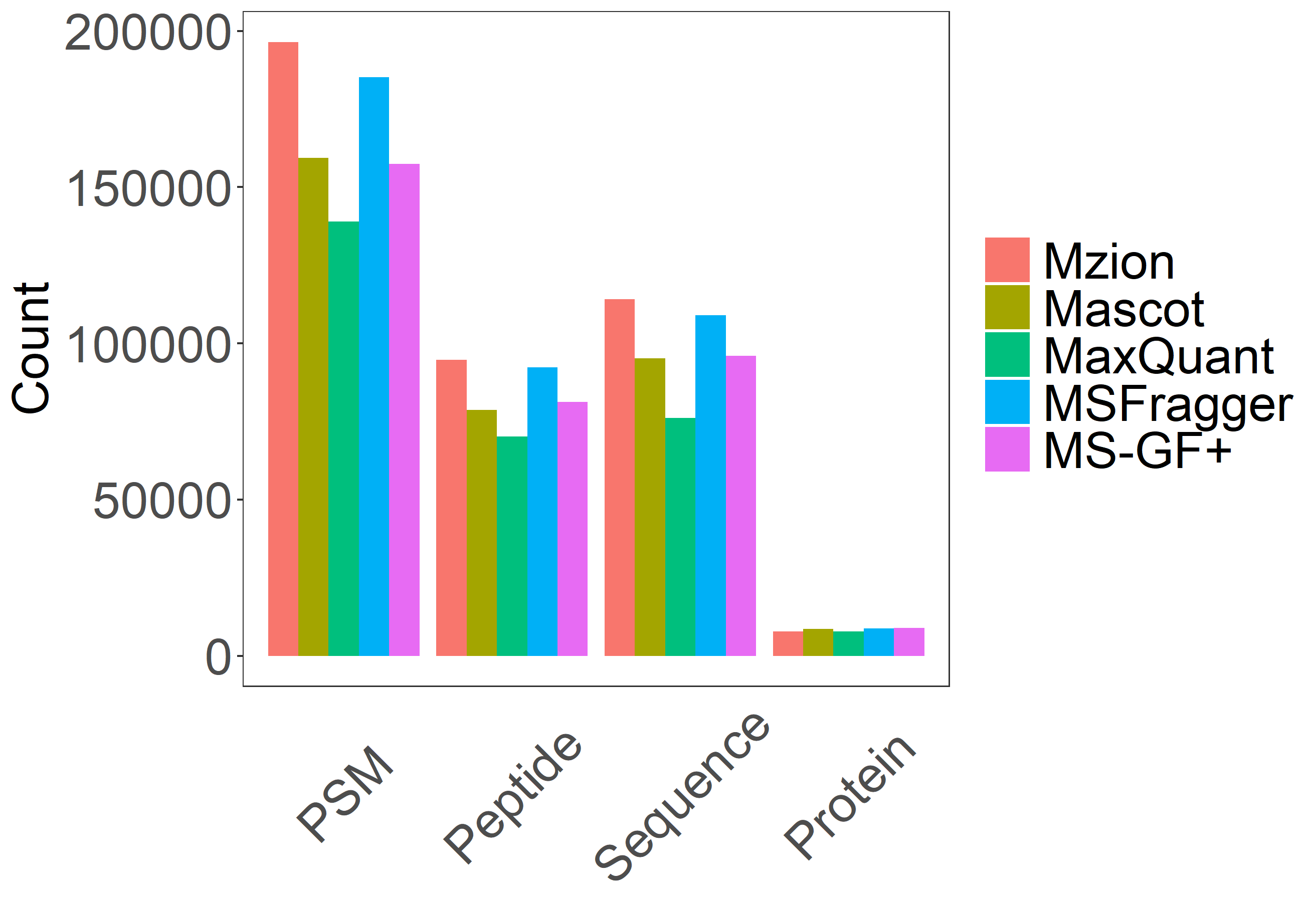

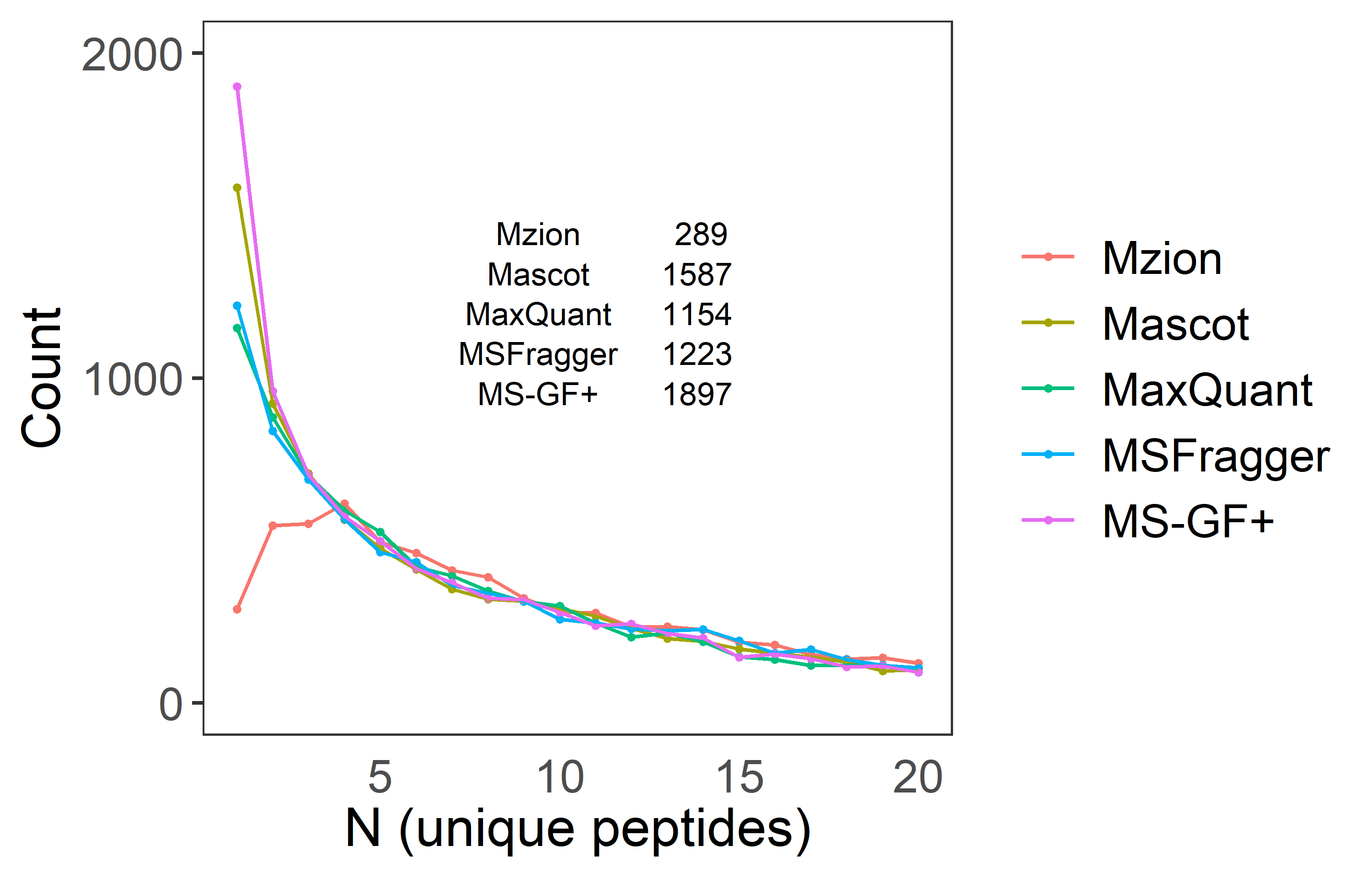

**b**

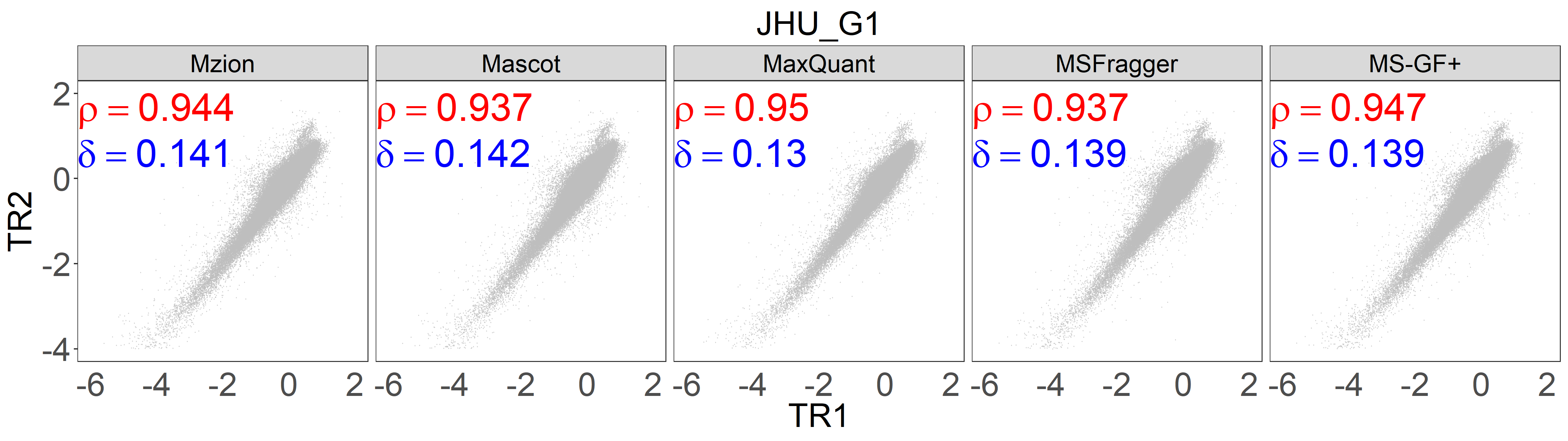

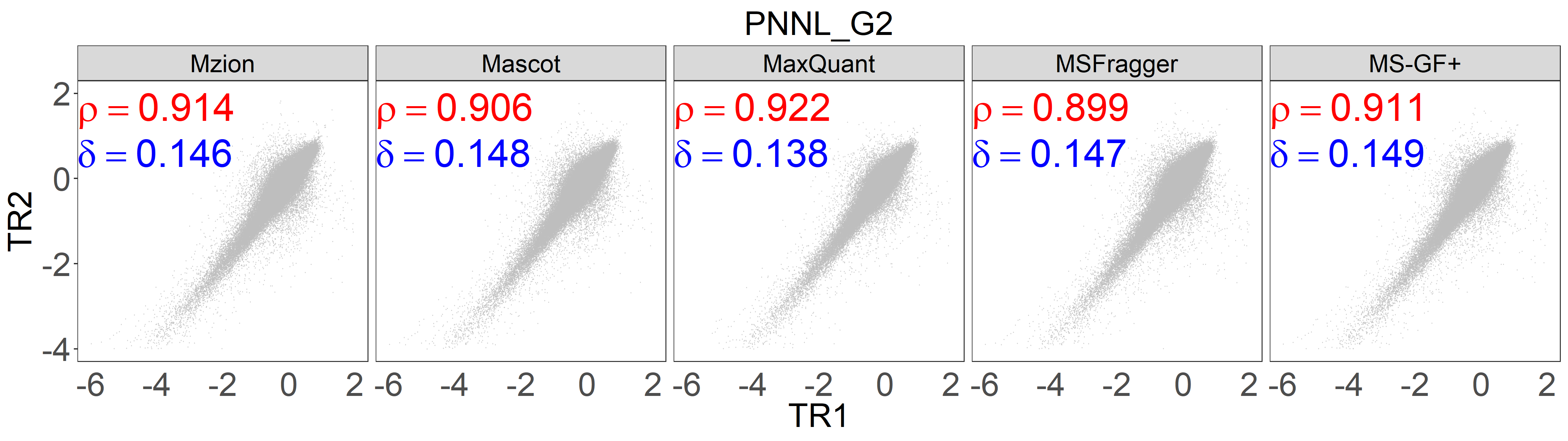

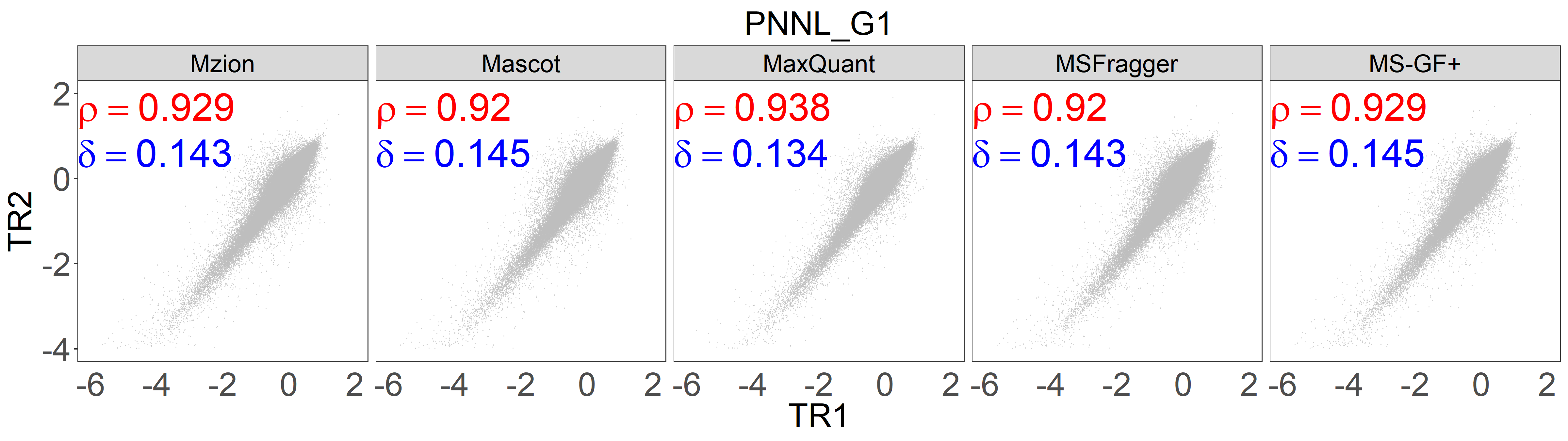

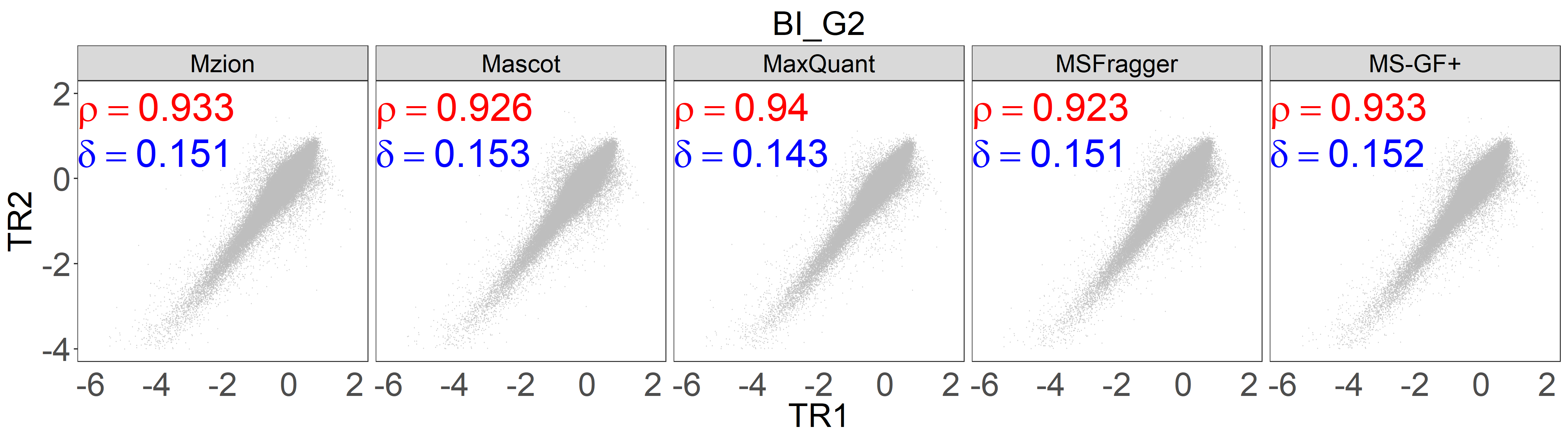
**Supplementary Figure 5.** Related to Figure 2. (a) Additional measures of Pearson correlations (red) and Manhattan distances (blue) of tryptic sequences between the first two WHIM2 replicates from dataset WHIM_G. The log2FC are relative to the average of five WHIM2 and five WHIM16 values. (b) Pairwise Manhattan distances of the WHIM2 replicates from dataset WHIM_G. (c) TMT0 entrapment against dataset WHIM_G. Peptides containing TMT0 modification on either K or peptide N-terminal are considered entrapped identities.

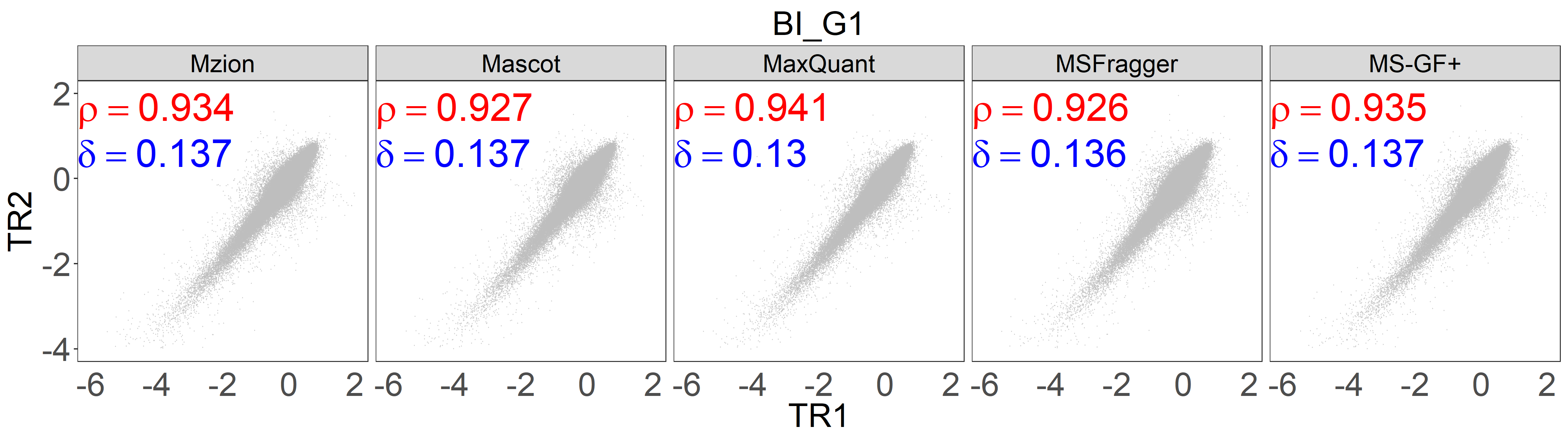

**a**

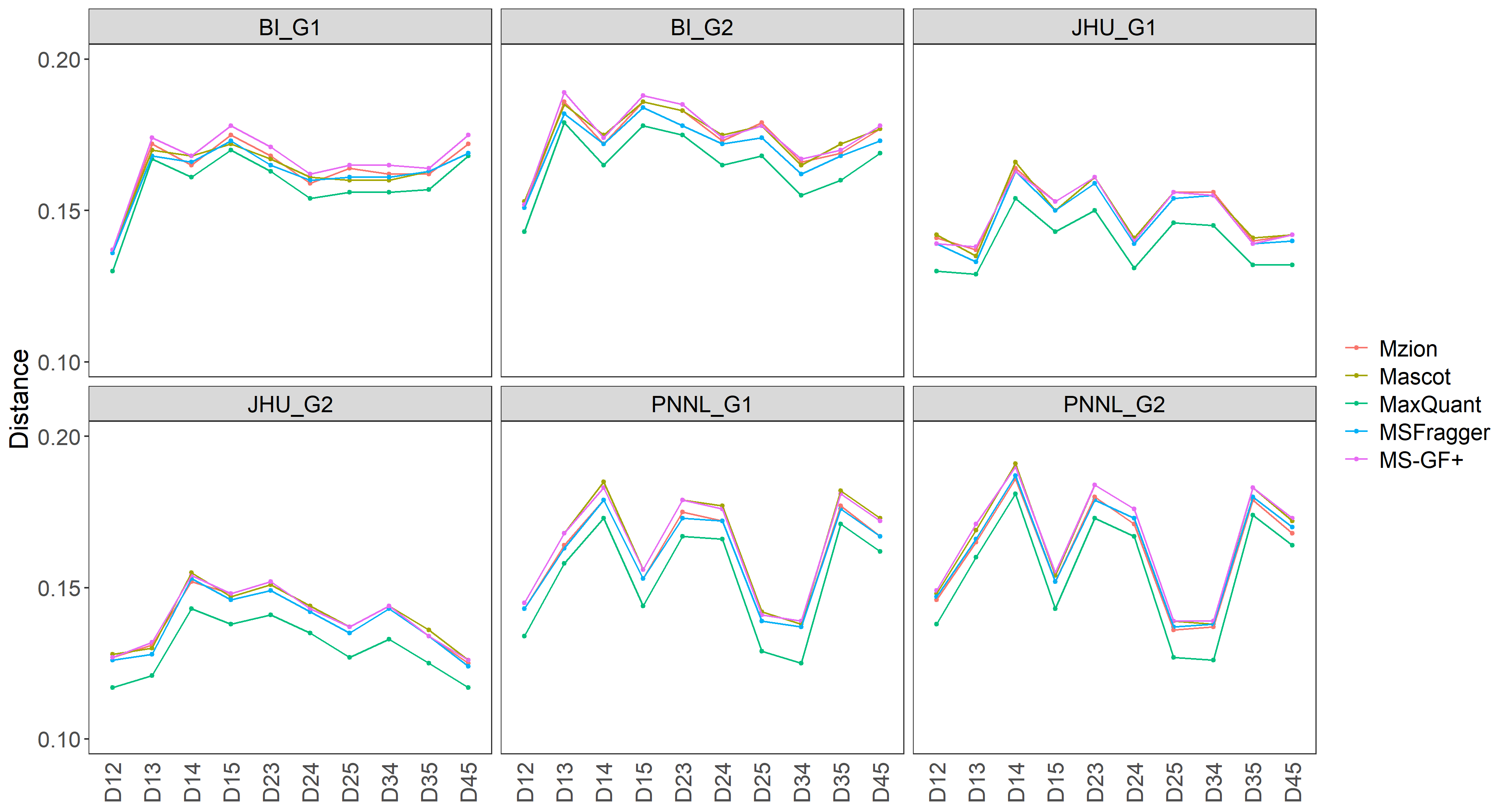

**b**

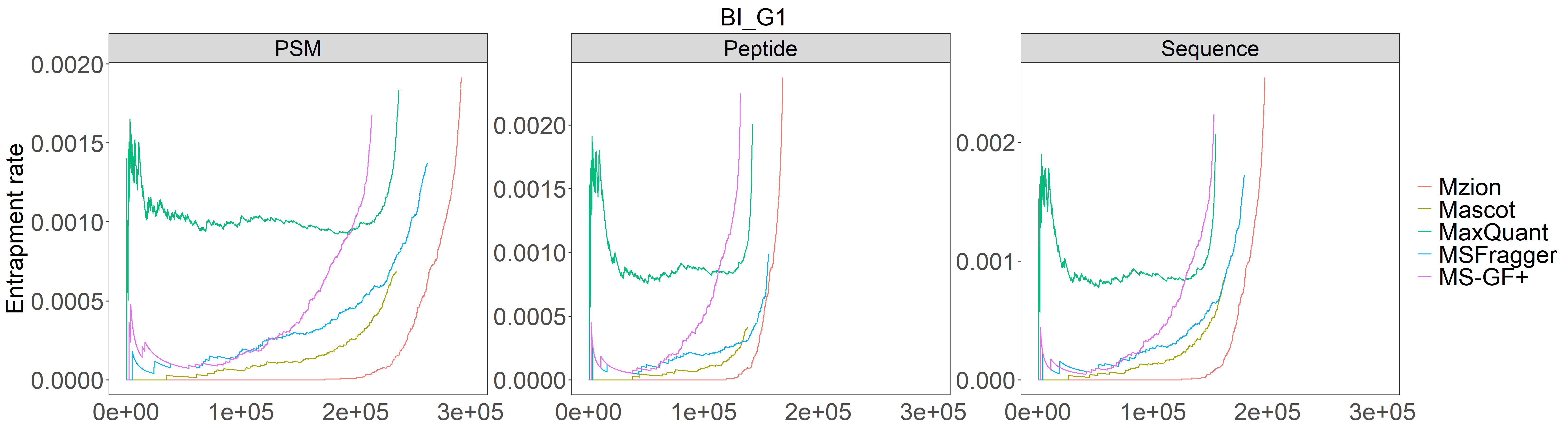

**c**

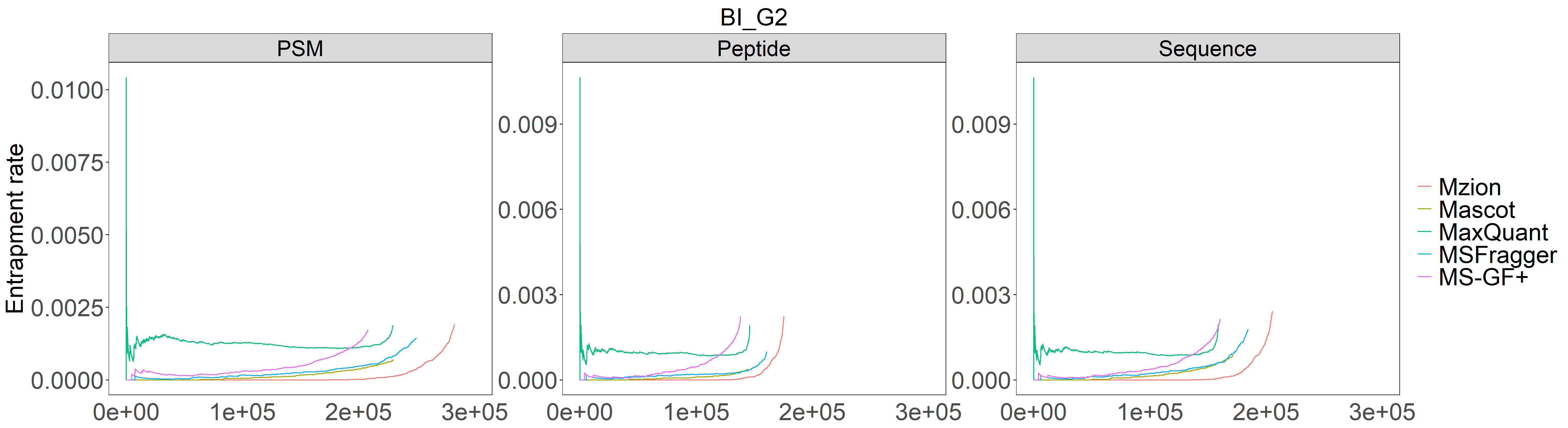

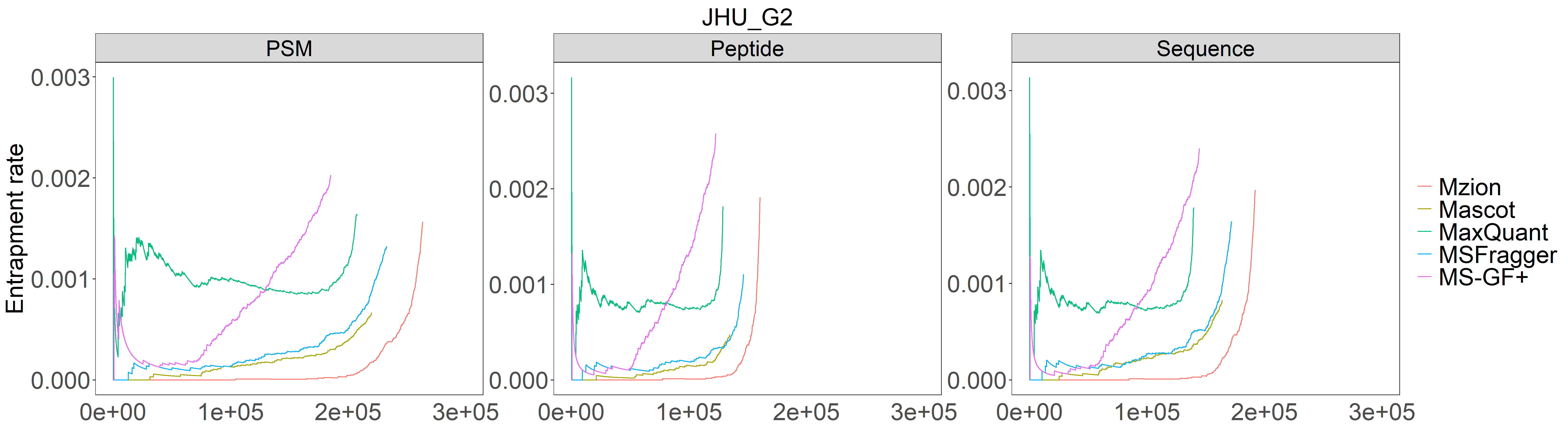

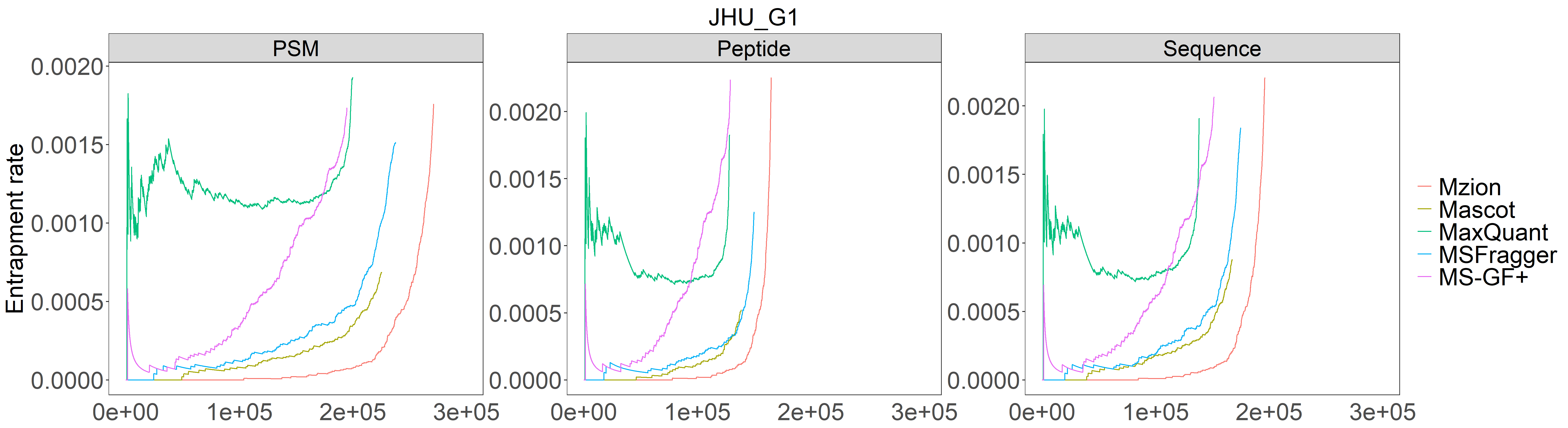

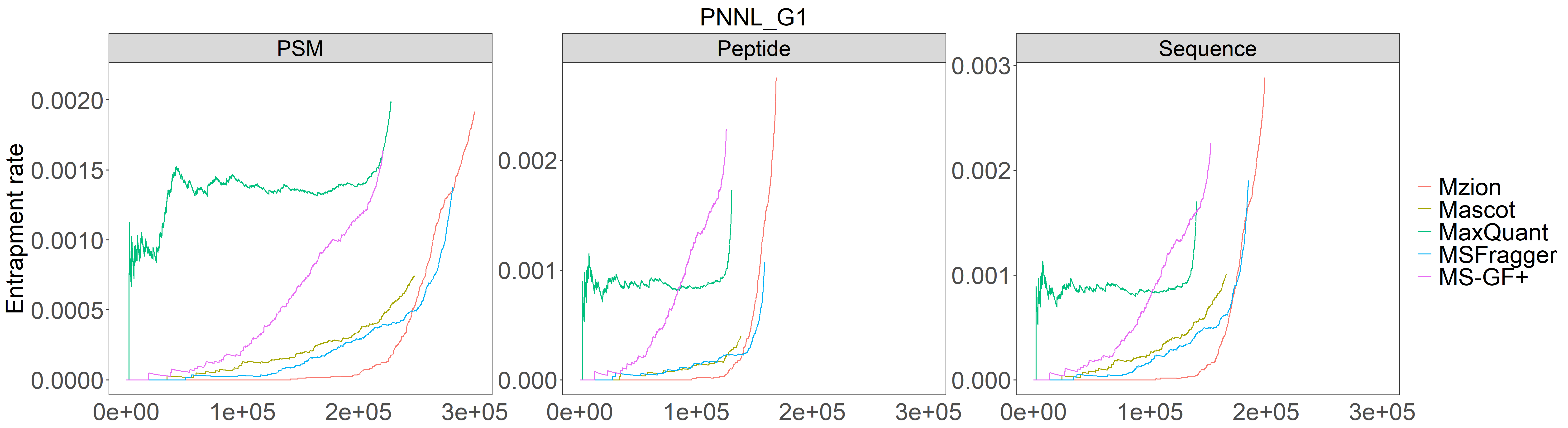

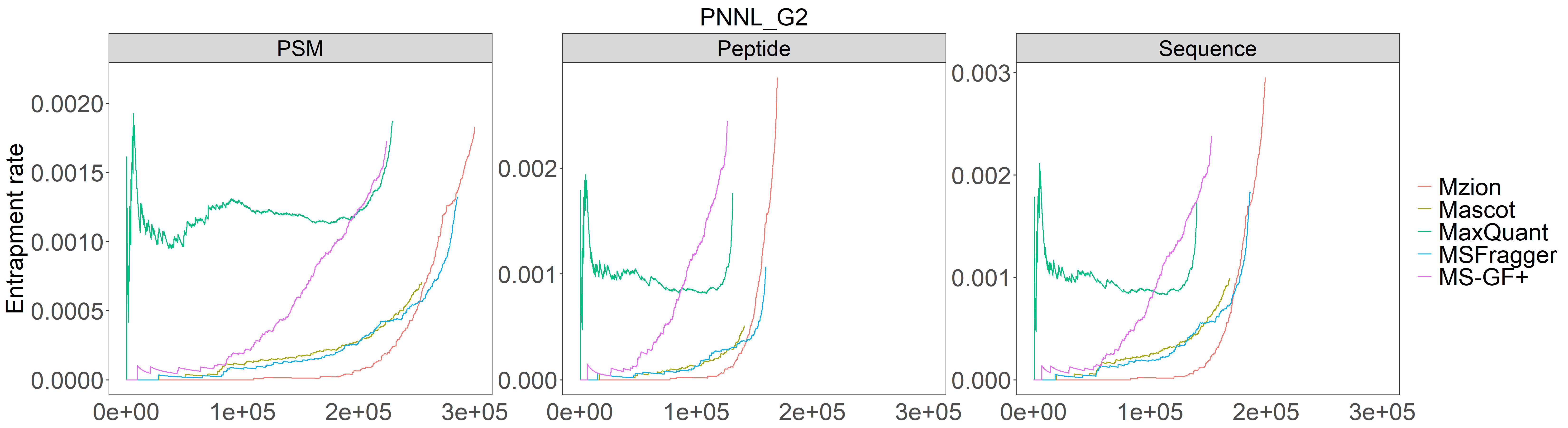

**Supplementary Figure 6.** Semi/non-tryptic findings from dataset WHIM_G.

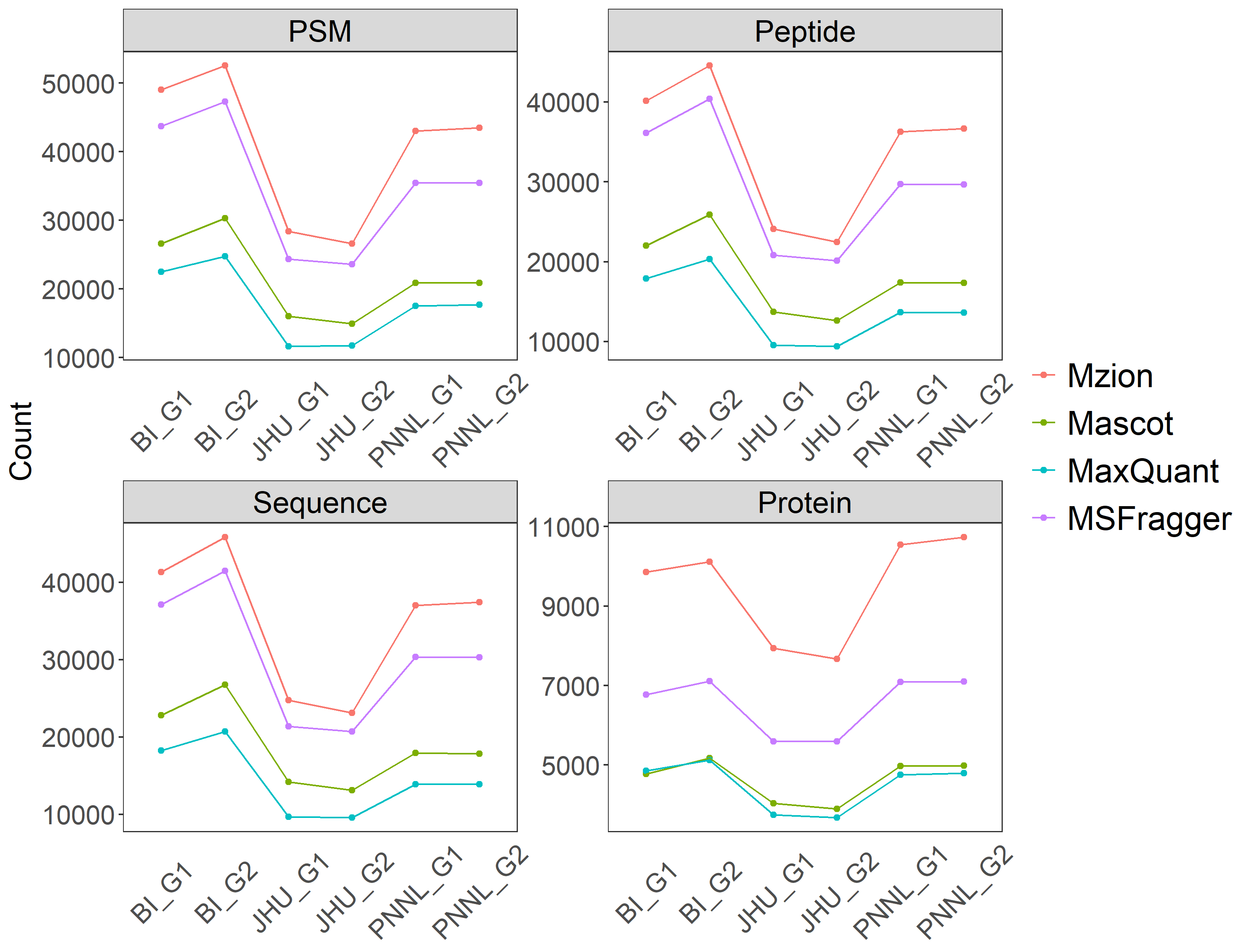

**Supplementary Figure 7.** Related to Figure 2. Additional measures of Pearson correlations (red) and Manhattan distances (blue) of semi/non-tryptic sequences between the first two WHIM2 replicates from dataset WHIM_G. NES: no enzyme specificity.

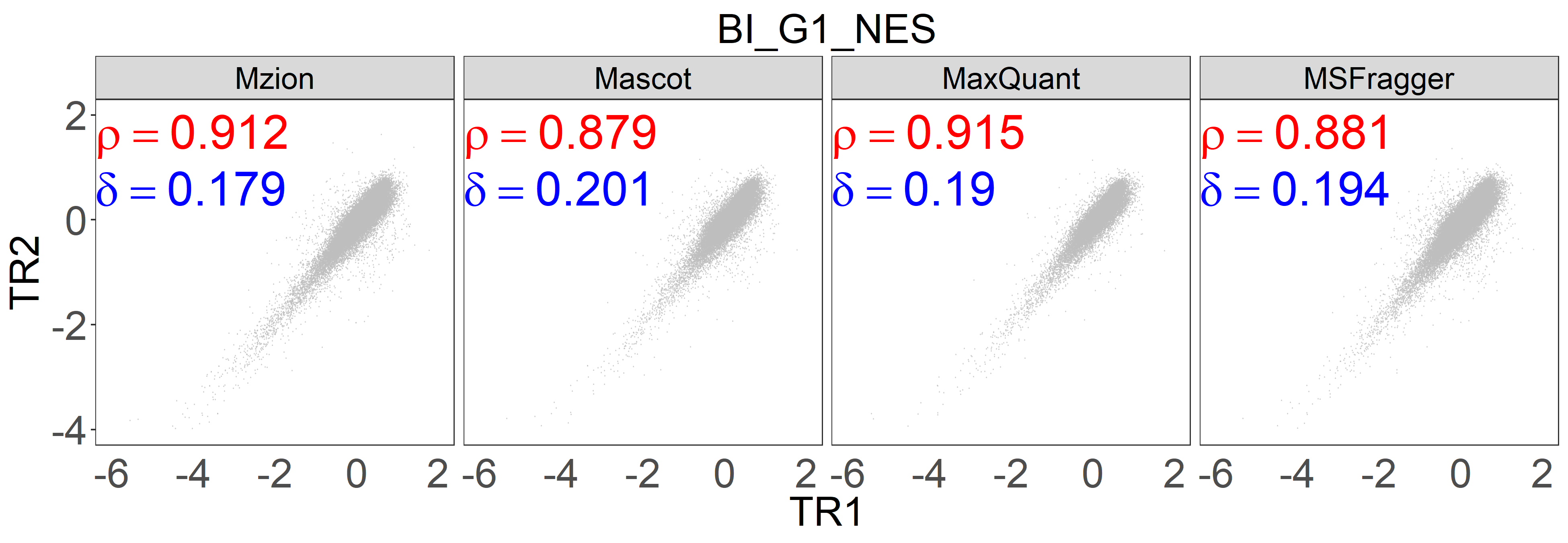

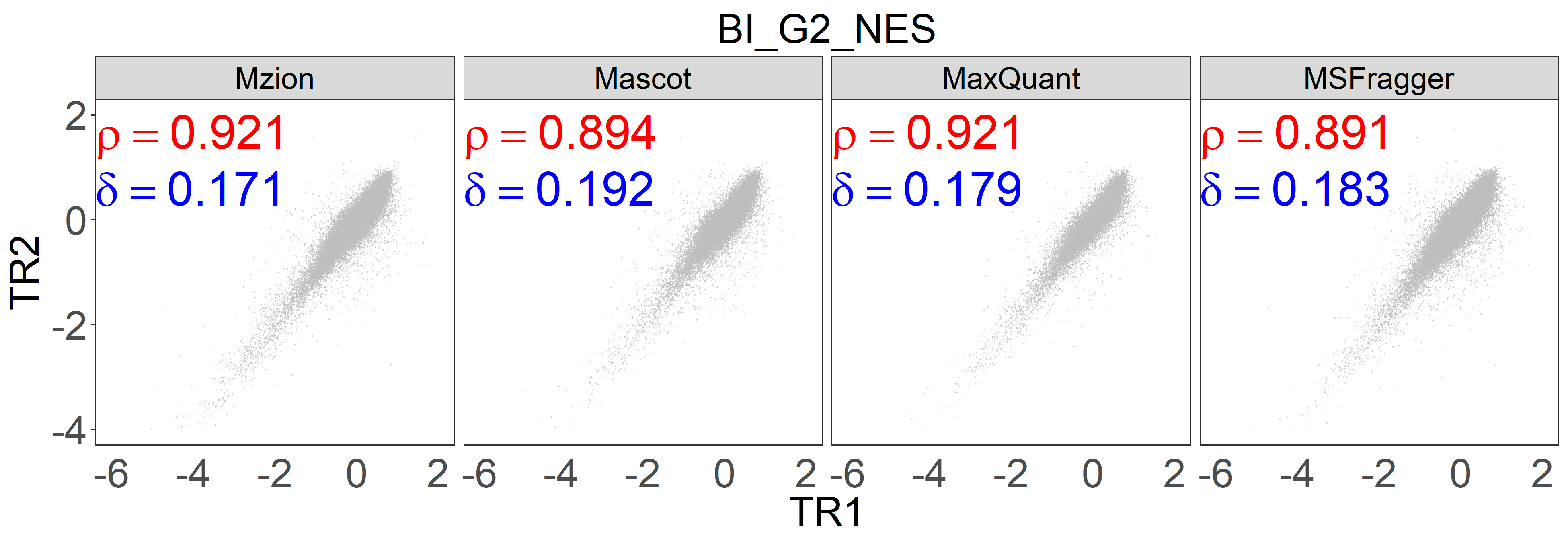

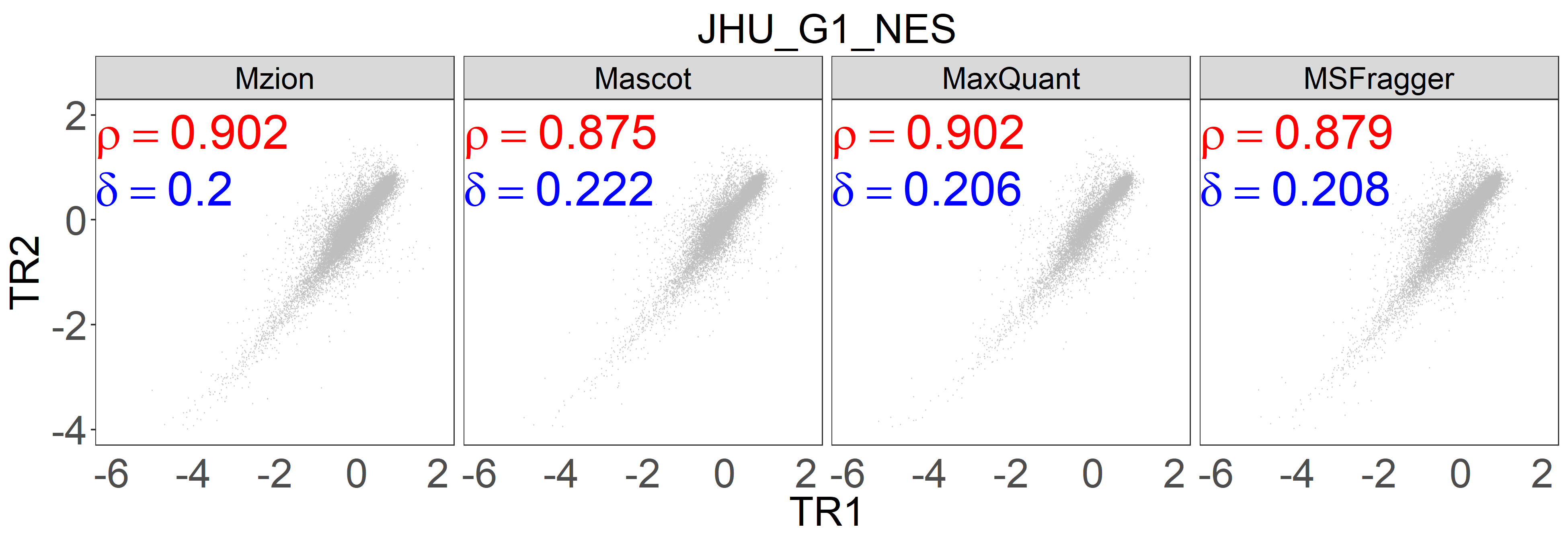

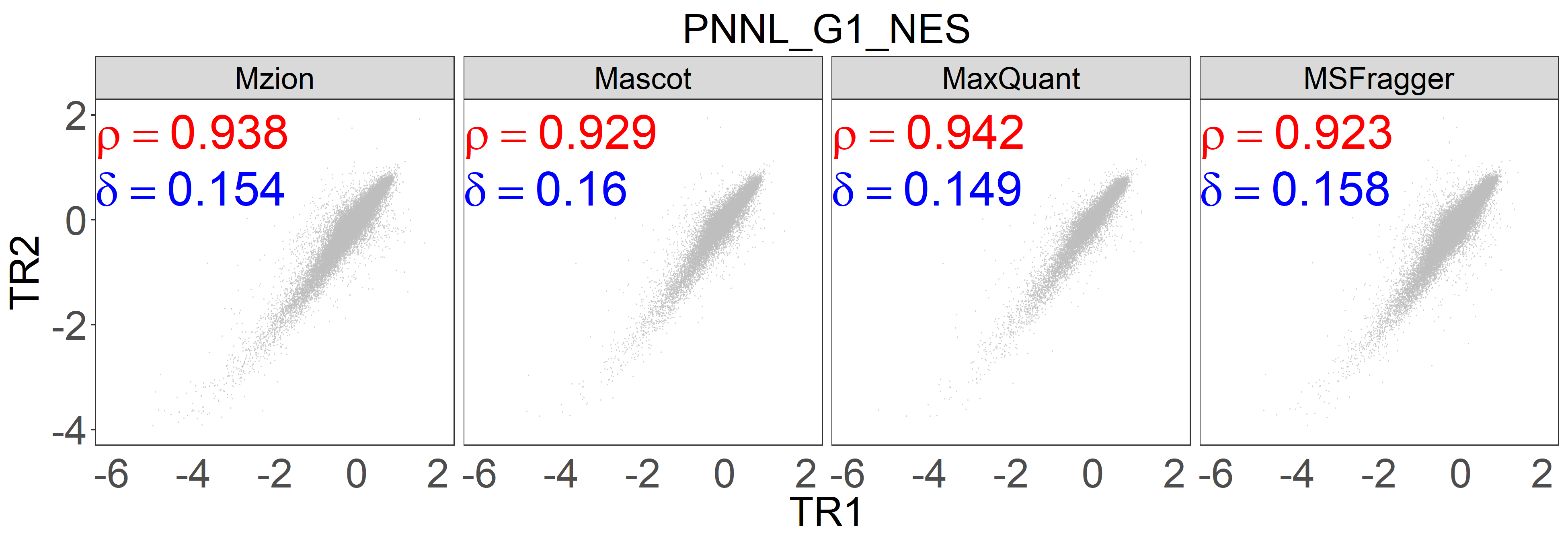

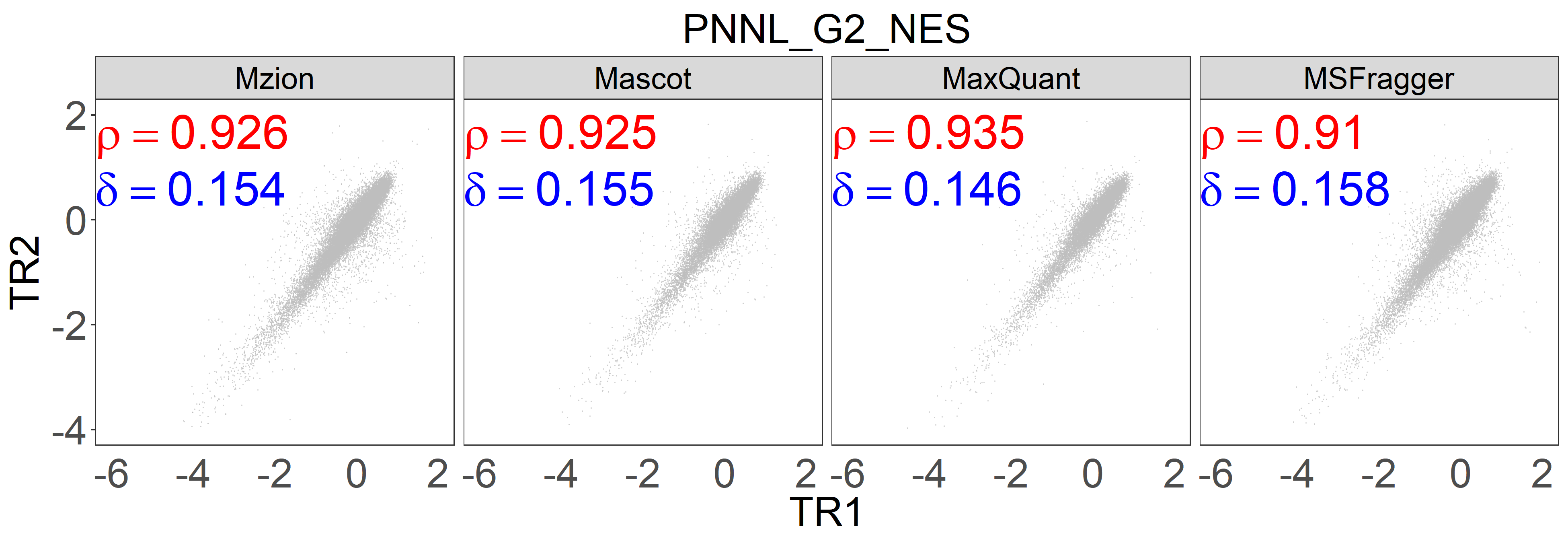

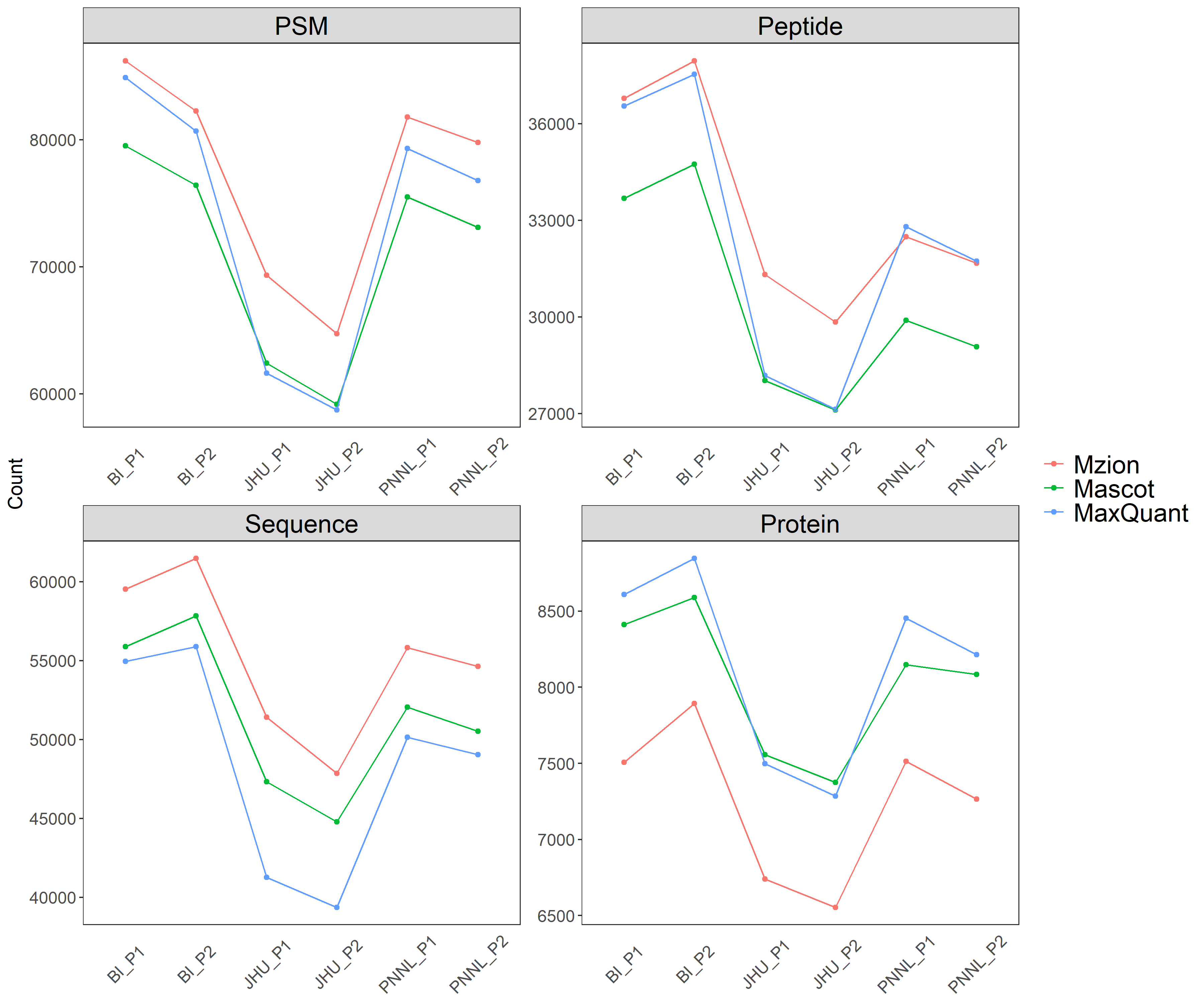
**Supplementary Figure 8.** Related to Figure 3. (a) Tryptic findings of from IMAC dataset WHIM_P. (b) Pairwise Manhattan distances of the WHIM2 replicates from IMAC dataset WHIM_P.

**a**

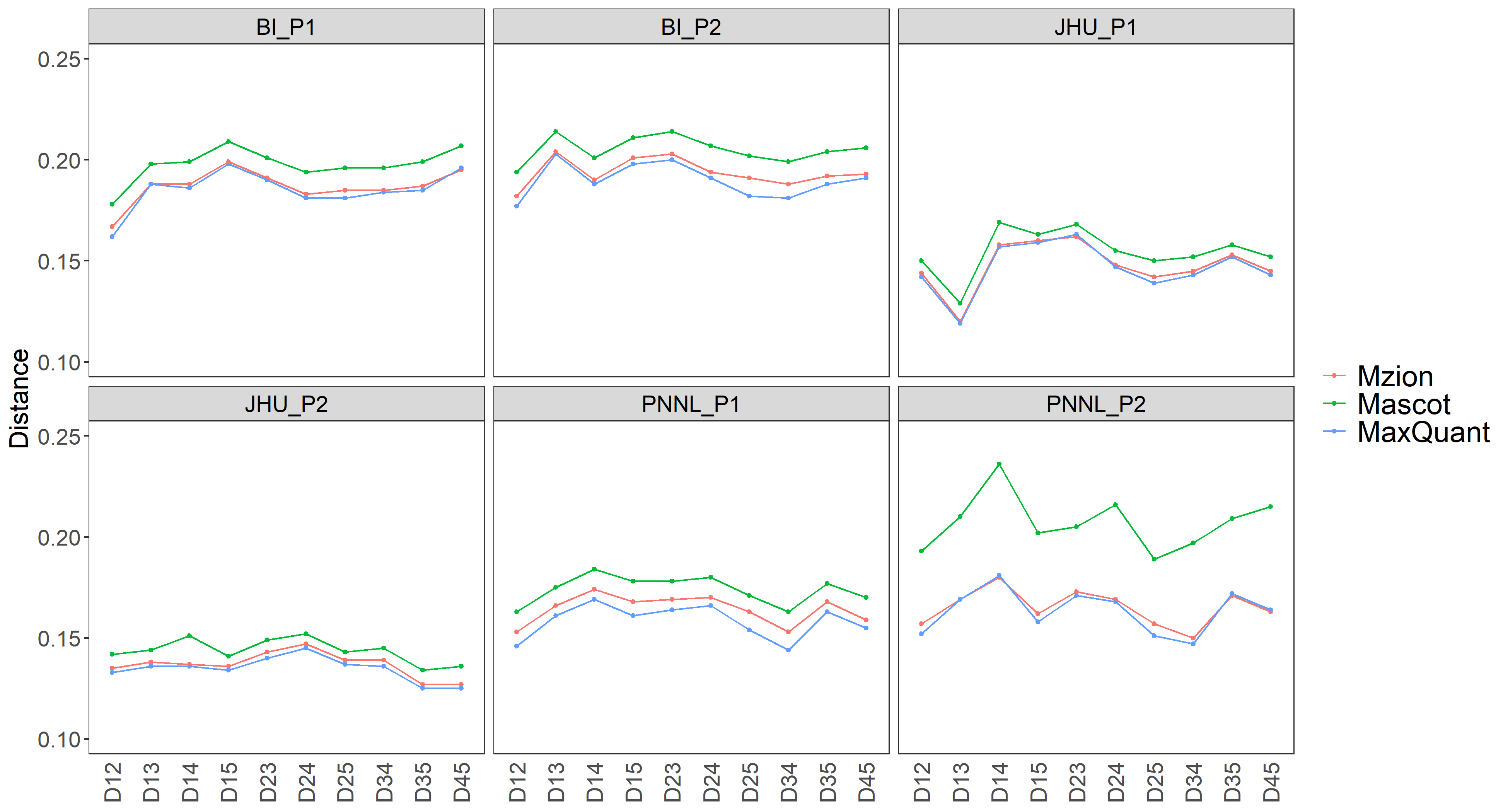

**b**

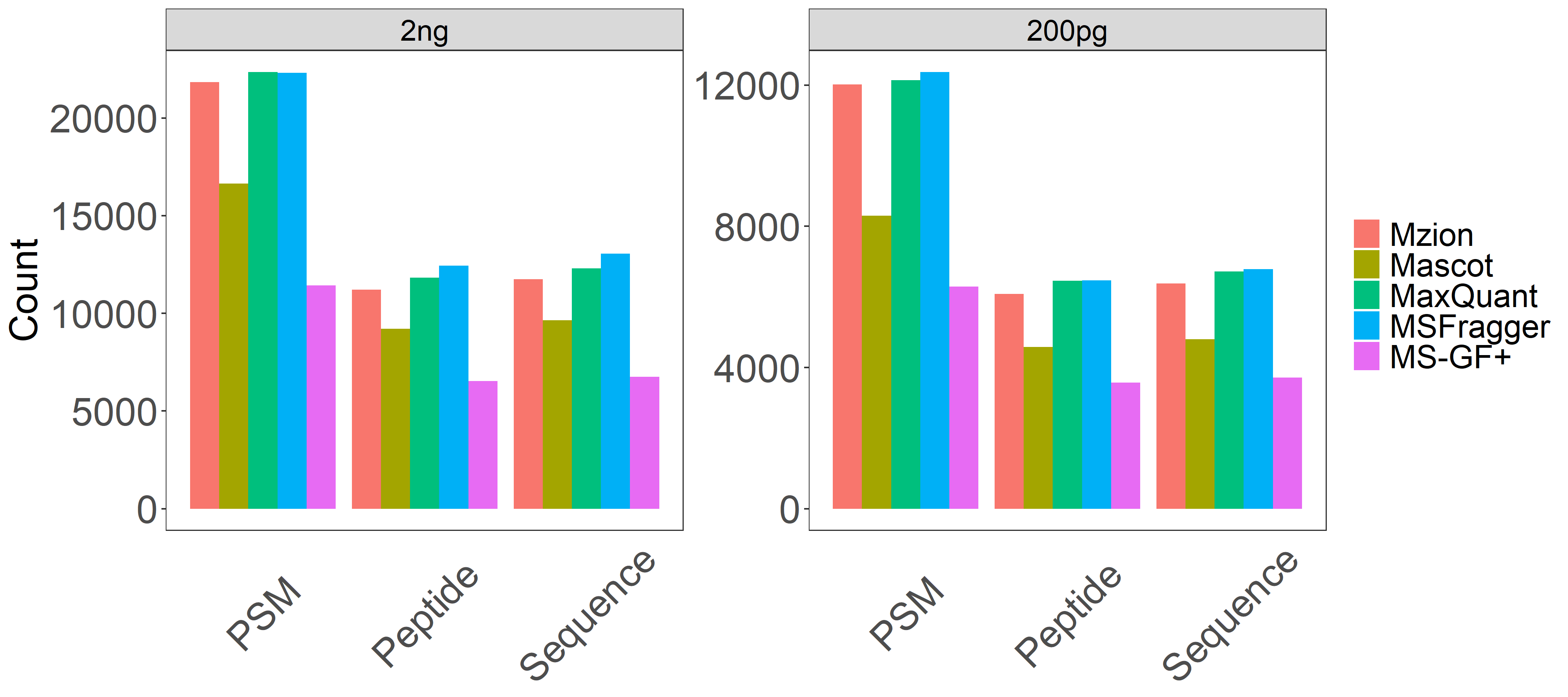
**Supplementary Figure 9.** Tryptic findings from datasets scHela_2ng and scHela_200pg. (**a**) Counts of PSMs, peptides and sequences. (**b**) Numbers of unique identifying peptides under proteins. Counts of proteins with one-peptide identification are shown in the inlet.

**a**

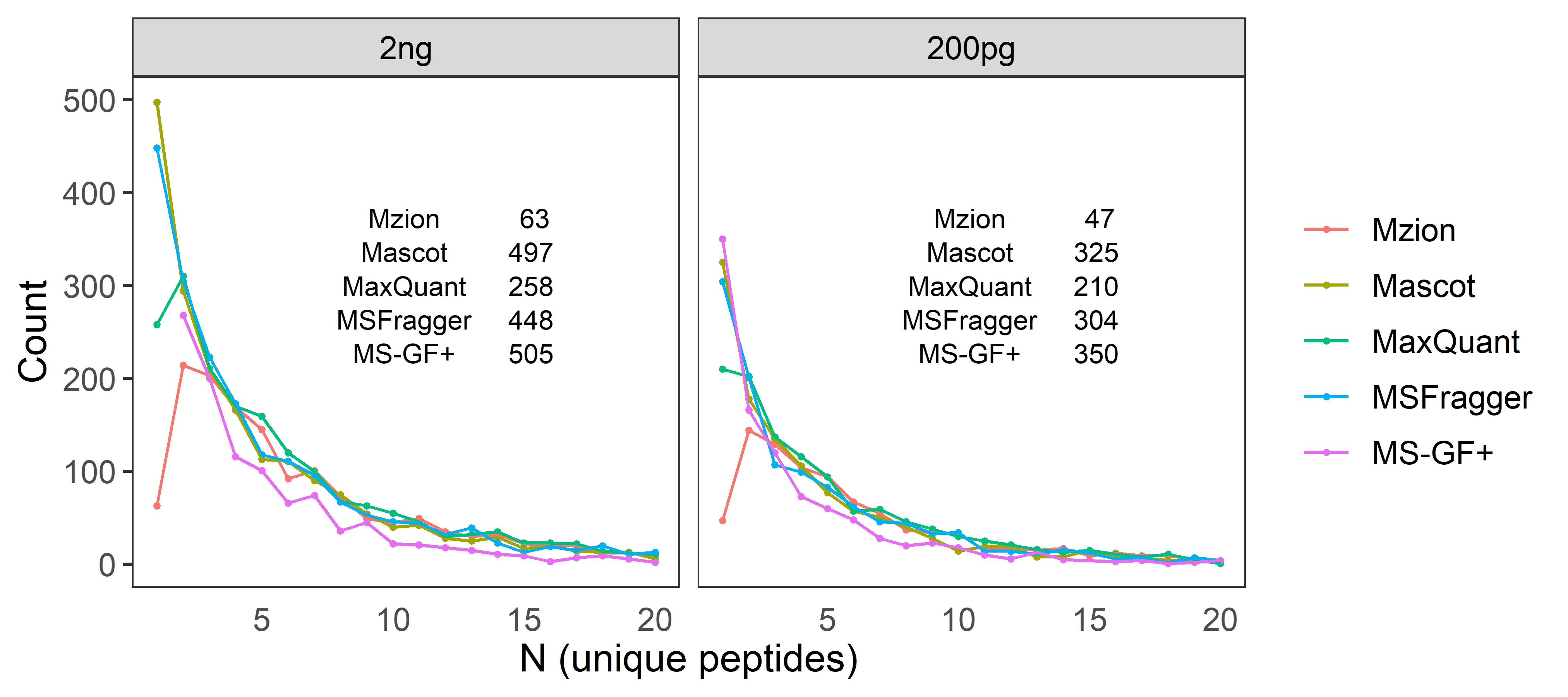

**b**

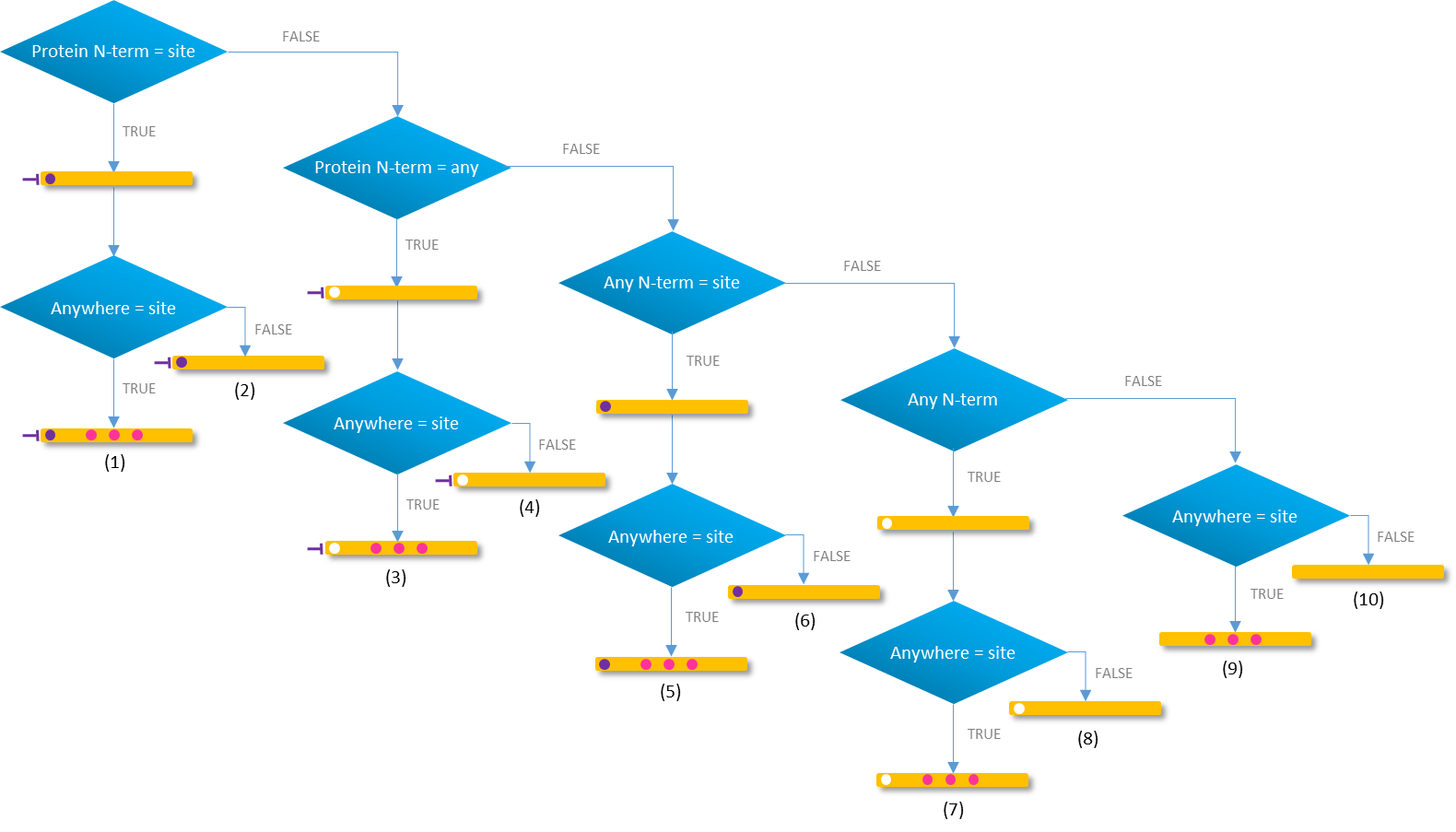

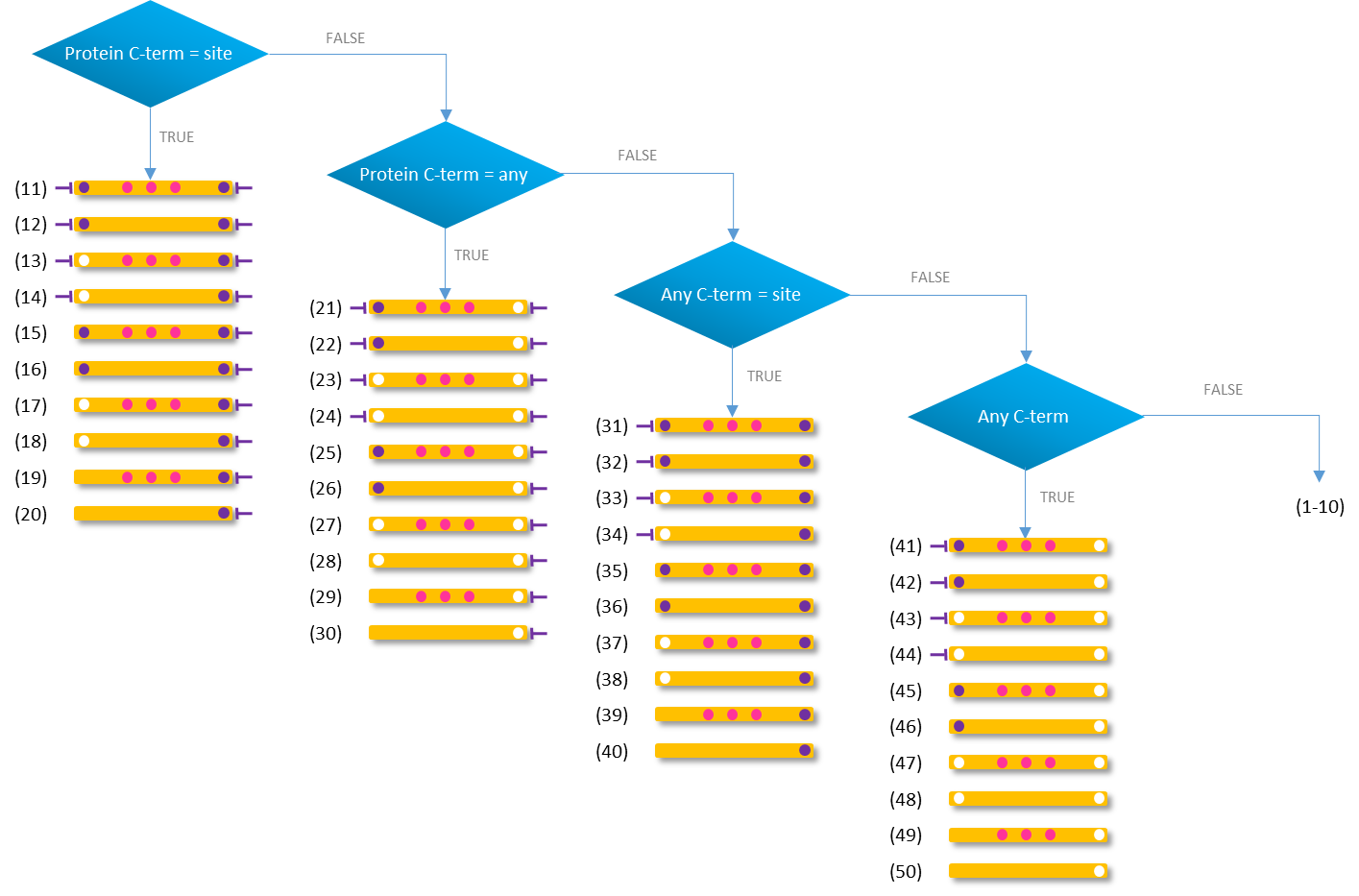
**Supplementary Figure 10.** Dispatching of precursor peptide sequences by variable modifications. The dispatching starts from the N-terminal tree, followed by the C-terminal tree.

Anywhere site

Protein N/C-term any site

Protein N/C-term and site

Protein C-term

Protein N-term

**Supplementary Figure 11.** Mzion findings from dataset BI_G1 at various search parameters. The gains from parameter set 1 to 2 or 3 are miniscule with the latter two coming at a cost of longer search time.

The searches were performed using Mzion v1.2.3. Common search parameters: database = Refseq human mouse (Jul. 2013; 56,673 entries) and cRAP (Jan. 2012, 116 entries); enzyme = trypsin/P, maximum missed cleavages = 4; minimum peptide length = 7; minimum precursor mass = 200, minimum product ion mass = 115, fixed modifications = cysteine carbamidomethylation, TMT10plex labeled lysine and TMT10plex labeled N-terminal; variable modifications = protein N-terminal acetylation, methionine oxidation, asparagine deamidation and N-terminal glutamine-to-pyroglutamic acid conversion; maximum variable modifications per peptide sequence = 5; maximum variable modifications per residue = 3; maximum combinations of variable modifications = 64; tolerance in precursor mass error = 20 ppm; offsets in 13C = 0; tolerance in product ion mass error = 20 ppm; tolerance in reporter-ion mass error = 10 ppm; FDR = 0.01 at PSMs, peptides and proteins.

Unique search parameters: (1) maximum peptide length = 40, maximum mass = 4500, top-N MS2 features = 100, minimum matched MS2 features = 6; (2) maximum peptide length = 50, maximum mass = 5000, top-N MS2 features = 100, minimum matched MS2 features = 6; (3) maximum peptide length = 50, maximum mass = 5000, top-N MS2 features = 150, minimum matched MS2 features = 6; (1) maximum peptide length = 50, maximum mass = 5000, top-N MS2 features = 150, minimum matched MS2 features = 5.

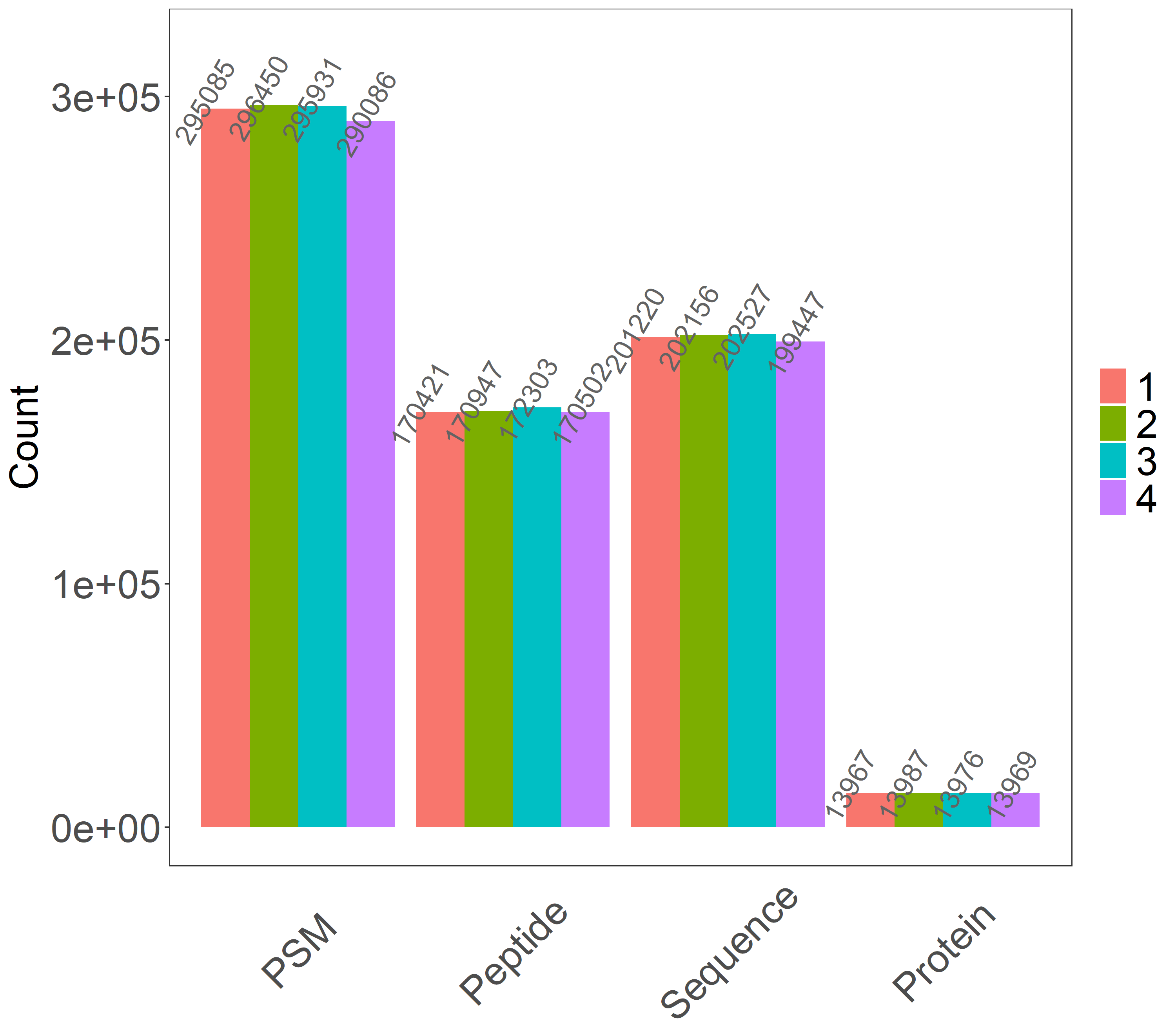
