## Supplementary Tables and Notes for "Mzion enables deep and precise identification of peptides in data-dependent acquisition proteomics"

**Supplementary Table 1.** Summary of modifications and abbreviations.

| Modification | Abbreviation |
| --- | --- |
| Asparagine deamidation | Deamidated (N) |
| Cysteine carbamidomethylation | Carbamidomethyl (C) |
| Lysine acetylation | Acetyl (K) |
| Lysine carbamylation | Carbamyl (K) |
| Lysine diglycination | Diglycyl (K) |
| Methionine oxidation | Oxidation (M) |
| N-terminal glutamine-to-pyroglutamic acid conversion | Gln->pyro-Glu (N-term = Q) |
| Protein N-terminal acetylation | Acetyl (Protein N-term) |
| TMT10plex labeled lysine | TMT10plex (K) |
| TMT10plex labeled lysine with diglycination | TMT10plex+Digly (K) |
| TMT10plex labeled N-terminal | TMT10plex (N-term) |
| TMTzero labeled lysine | TMTzero (K) |
| TMTzero labeled N-terminal | TMTzero (N-term) |

**Supplementary Table 2.** Sets of fixed and variable modifications at a user’s specification of fixed Carbamidomethyl (C), TMT10plex (N-term) and TMT10plex (K) and variable Acetyl (Protein N-term), Gln->pyro-Glu (N-term = Q), Oxidation (M) and Deamidated (N). A principle of minimalism is employed when compiling the final set of modifications. For instance, modifications Carbamidomethyl (C) and TMT10plex (K) both remain as fixed when there are no other variable modifications to sites C and K. Assumption of fixed TMT10plex (N-term) is lifted when a variable N-terminal modification is present. Combinations without an N-terminal modification are removed given the user’s intention of a fixed N-terminal modification. An exemplary breakdown of total PSM identifications from dataset BI_G1 showed that the top-4 modification sets are associated with the fixed TMT10plex (N-term) modification.

| Group | Fixed | Variable | PSM |
| --- | --- | --- | --- |
| 1 | Carbamidomethyl (C), TMT6plex (K), TMT6plex (N-term) |  | 238189 |
| 2 | Carbamidomethyl (C), TMT6plex (K) | Acetyl (Protein N-term) | 3076 |
| 3 | Carbamidomethyl (C), TMT6plex (K) | Gln->pyro-Glu (N-term = Q) | 1762 |
| 4 | Carbamidomethyl (C), TMT6plex (K), TMT6plex (N-term) | Deamidated (N) | 24934 |
| 5 | Carbamidomethyl (C), TMT6plex (K), TMT6plex (N-term) | Oxidation (M) | 31524 |
| 6 | Carbamidomethyl (C), TMT6plex (K) | Acetyl (Protein N-term), Deamidated (N) | 321 |
| 7 | Carbamidomethyl (C), TMT6plex (K) | Acetyl (Protein N-term), Oxidation (M) | 817 |
| 8 | Carbamidomethyl (C), TMT6plex (K) | Deamidated (N), Gln->pyro-Glu (N-term = Q) | 129 |
| 9 | Carbamidomethyl (C), TMT6plex (K) | Gln->pyro-Glu (N-term = Q), Oxidation (M) | 73 |
| 10 | Carbamidomethyl (C), TMT6plex (K), TMT6plex (N-term) | Deamidated (N), Oxidation (M) | 4685 |
| 11 | Carbamidomethyl (C), TMT6plex (K) | Acetyl (Protein N-term), Deamidated (N), Oxidation (M) | 185 |
| 12 | Carbamidomethyl (C), TMT6plex (K) | Deamidated (N), Gln->pyro-Glu (N-term = Q), Oxidation (M) | 5 |

**Supplementary Table 3.** Summary of twenty datasets from six studies. The six global datasets from reference Mertins et al. are termed collectively WHIM_G and the six IMAC-enriched phosphopeptide datasets termed WHIM_P.

| Dataset | Type | ID | Ref |
| --- | --- | --- | --- |
| BI_G1 | Global | PDC000311 | Mertins et al. (2018)^1^ |
| BI_G2 |  |  |  |
| JHU_G1 |  |  |  |
| JHU_G2 |  |  |  |
| PNNL_G1 |  |  |  |
| PNNL_G2 |  |  |  |
| BI_P1 | IMAC | PDC000314 |  |
| BI_P2 |  |  |  |
| JHU_P1 |  |  |  |
| JHU_P2 |  |  |  |
| PNNL_P1 |  |  |  |
| PNNL_P2 |  |  |  |
| Pancr_G1 | Global | PDC000270-1 | Cao et al. (2021)^2^ |
| Pancr_P1 | IMAC | PDC000271-1 |  |
| Ferris_spike | IMAC | PXD007058-HCDOT | Ferries et al. (2017)^3^ |
| Lung_A1_A10 | Acetyl | PDC000233 | Satpathy et al. (2021)^4^ |
| Lung_U | Ubiquintyl | PDC000237 |  |
| Dong_SILAC | SILAC | Dong-Ecoli-QE | Chi et al. (2018)^5^ |
| scHela_200pg | Single-cell | MSV000087524 | Boekweg et al. (2022)^6^ |
| scHela_2ng |  |  |  |

| **Supplementary Table 4.** Semi/non-tryptic findings from the global dataset Panc_G1. | | | | | | | | | | | | | | | |
| --- | --- | --- | --- | --- | --- | --- | --- | --- | --- | --- | --- | --- | --- | --- | --- |
|  | | | |  | PSM | |  | Peptide | |  | Sequence | |  | Protein | |
| Engine | PSM | Peptide | Sequence | Protein | ! Q% | ! C% |  | ! Q% | ! C% |  | ! Q% | ! C% |  | ! Q% | ! C% |
| Mzion | 77,853 | 54,608 | 57,044 | 7,580 |  |  |  |  |  |  |  |  |  |  |  |
| Mascot | 48,878 | 32,488 | 34,439 | 4,565 | 7.2 | 0.0 |  | 6.1 | 0.3 |  | 6.2 | 0.3 |  | 3.6 | 0.1 |
| MaxQuant | 31,467 | 20,737 | 21,347 | 3,731 | 10.3 | 0.5 |  | 4.9 | 0.3 |  | 37.4 | 5.0 |  | 2.3 | 0.1 |
| MSFragger | 59,935 | 39,839 | 42,084 | 5,151 | 15.1 | 0.1 |  | 12.6 | 0.5 |  | 13.1 | 0.5 |  | 4.8 | 0.1 |
| ^1^ ! Q% - Percent not in the quality space of psmQ.txt | | | | | | | | | | | | | | | |
| ^2^ ! C% - Percent not in the complete space of psmC.txt | | | | | | | | | | | | | | | |

| **Supplementary Table 5.** Findings of spiked phosphopeptides from dataset Ferries_spikes. | | | | | | | | | | | |
| --- | --- | --- | --- | --- | --- | --- | --- | --- | --- | --- | --- |
|  | | | | PSM | |  | Peptide | |  | Sequence | |
| Engine | PSM | Peptide | Sequence | ! Q% | ! C% |  | ! Q% | ! C% |  | ! Q% | ! C% |
| Mzion | 797 | 98 | 180 |  |  |  |  |  |  |  |  |
| Mascot | 894 | 117 | 212 | 26.9 | 0.1 |  | 21.4 | 0.0 |  | 32.2 | 1.2 |
| MaxQuant | 853 | 145 | 230 | 40.9 | 2.1 |  | 54.1 | 1.1 |  | 53.9 | 2.5 |
| MSFragger | 965 | 122 | 256 | 40.8 | 0.2 |  | 28.6 | 0.1 |  | 80.6 | 4.8 |
| MS-GF+ | 800 | 112 | 228 | 56.2 | 0.3 |  | 31.6 | 0.4 |  | 76.1 | 1.7 |
| ^1^ ! Q% - Percent not in the quality space of psmQ.txt | | | | | | | | | | | |
| ^2^ ! C% - Percent not in the complete space of psmC.txt | | | | | | | | | | | |

**Supplementary Table 6.** Tryptic findings from the SILAC dataset Dong_ecoli.

| Set | Engine | PSM | Peptide | Sequence | Protein |  | PSM | |  | Peptide | |  | Sequence | |  | Protein | |
| --- | --- | --- | --- | --- | --- | --- | --- | --- | --- | --- | --- | --- | --- | --- | --- | --- | --- |
|  |  |  |  |  |  |  | !Q | !C |  | !Q | !C |  | !Q | !C |  | !Q | !C |
| base | Mzion | 36,309 | 14,023 | 16,319 | 1,724 |  |  |  |  |  |  |  |  |  |  |  |  |
|  | Mascot | 29,011 | 11,874 | 13,934 | 1,835 |  | 1.5 | 0.9 |  | 1.7 | 0.6 |  | 4.1 | 2.8 |  | 7.7 | 0.3 |
|  | MSFragger | 21,402 | 10,241 | 11,933 | 1,764 |  | 2.5 | 1.2 |  | 2.4 | 0.7 |  | 7.1 | 5.1 |  | 9.3 | 0.8 |
| grpC | Mzion | 22,575 | 10,767 | 11,851 | 1,437 |  |  |  |  |  |  |  |  |  |  |  |  |
|  | Mascot | 15,697 | 8,870 | 9,043 | 1,679 |  | 2.9 | 1.3 |  | 4.0 | 1.3 |  | 5.4 | 2.8 |  | 18.2 | 0.3 |
|  | MSFragger | 17,862 | 9,085 | 10,035 | 1,686 |  | 4.9 | 1.8 |  | 5.0 | 1.3 |  | 8.4 | 4.6 |  | 19.9 | 1.0 |
| grpN | Mzion | 27,954 | 12,223 | 13,583 | 1,595 |  |  |  |  |  |  |  |  |  |  |  |  |
|  | Mascot | 18,976 | 9,439 | 9,619 | 1,681 |  | 1.2 | 0.4 |  | 1.6 | 0.4 |  | 4.0 | 2.7 |  | 7.7 | 0.5 |
|  | MSFragger | 23,137 | 10,690 | 11,798 | 1,812 |  | 8.3 | 6.1 |  | 5.9 | 3.1 |  | 10.5 | 7.6 |  | 15.5 | 1.8 |
| ^1^ ! Q% - Percent not in the quality space of psmQ.txt  ^2^ ! C% - Percent not in the complete space of psmC.txt  ^3^ base – Sample group without stable isotope incorporation | | | | | | | | | | | | | | | | | |
| ^4^ grpC – Sample group with metabolic incorporation of ^13^C | | | | | | | | | | | | | | | | | |
| ^5^ grpN – Sample group with metabolic incorporation of ^15^N | | | | | | | | | | | | | | | | | |

Supplementary Notes

### Supplementary Note 1

#### Handling the competing specifications between fixed and variable modifications.

It is a common exercise in proteomics publications to provide both fixed and variable modifications that have been assumed in database searches. A variable modification can be realized permutatively on one or multiple sites in a peptide sequence. Under simple circumstances, the mass of a realized variable modification (variable mass) can be applied additively to that of a fixed modification (fixed mass) at the same site of a peptide sequence. However, additions or subtractions of variable masses ignoring chemistry are inappropriate and were often handled furthermore by search engines.

To illustrate, we suppose an experiment considering tentatively the fixed modifications of carbamidomethylation of cystein (Carbamidomethyl (C)), TMT-10plex labeling of lysine (TMT10plex (K)) and TMT-10plex labeling of peptide N-terminals (TMT10plex (N-term)), and the variable modifications of protein N-terminal acetylation (Acetyl (Protein N-term)) and N-terminal glutamine-to-pyroglutamic acid conversion (Gln->pyro-Glu (N-term)). At the combination of variable Acetyl (Protein N-term) and fixed TMT10plex (N-term), adding the variable mass to the fixed mass is chemically unsound. Probably for this reason, Mascot turns off the specification of fixed TMT10plex (N-term) when encountering a protein N-terminal acetylation. While it is suitable to escape from competing modifications by special case handling, the approach cannot be generalized without knowing apriori the choices by experimenters.

Another resort is to expand the search space by making TMT10plex (N-term) a variable modification, an approach appears to be taken implicitly by MaxQuant (internally programmed and cannot be modified by users) or explicitly by Mascot (e.g. at a quantitation method of “TMT10plex (variable)”). The arrangement introduces extraneously the combination of free N-terminal (without TMT10plex) to fixed Carbamidomethyl (C), as well as the combinations of free N-terminal to every other variable modification. Note that tryptic peptides without TMT10plex on either residue K or N-terminal will have no quantitative values but remain in the search space. It then becomes users’ responsibility to remove the corresponding entries from quantitation, which may not be a trivial task. With the approach of search space expansion, software(s) may have a higher propensity in ascribing inadvertently reporter-ion signals from co-isolated and co-fragmented species^7,8^ to peptide sequences that contains no TMT10plex, resulting in superfluous protein and peptide quantitations. Taking an example of MS data 01CPTAC3_Benchmarking_W_BI_20170508_BL_f19 at scan 47933 and precursor mass 1936.8637 from dataset BI_G, it may have been reported as a high confidence match to deamidated AIEDSDMLQETMEEYMNKPTF with descent reporter-ion intensities. However, provided the precursor mass, the match to AIEDSDMLQETMEEYMNKPTF should contain no TMT10plex modifications on either peptide N-terminal or K; hence the reporter-ion signals do not belong to the ascribed peptide sequence. Indeed, mismatches of reporter-ion intensities to peptide sequences without TMT labels were widely observed with search engines.

Search engines such as MSFragger and MS-GF+ are capable of addressing the problem with mass subtractions. For instance, the fixed mass of TMT10plex (N-term) is first applied universally. The variable mass of Acetyl (Protein N-term) is then supplied as a delta between the mass of TMT10plex (N-term) and the mass of the original Acetyl (Protein N-term). The approach works well but comes at a cost of users’ responsibility in deducing the compatibility between competing fixed and variable modifications. The task might become hefty when the combinations of modifications are complex.

Mzion attempts to reconcile the competing fixed and variable modifications in a more general way and to alleviate the users’ responsibility in deducing the compatibility between the two types of modifications. The principle of parsimony is applied to reduce excessive expansions of search space. In the example of user-specified fixed Carbamidomethyl (C), TMT10plex (K) and TMT10plex (N-term), and variable Acetyl (Protein N-term), Gln->pyro-Glu (N-term = Q), Deamidated (N) and Oxidation (M), Mzion results in a set of twelve combinations of fixed and variable modification (Supplementary Table 2). The search engine first generates all possible combinations and excludes those without the site of competition, which is N-terminal in the case. The user-defined fixed modification, which is TMT10plex (N-term) in the example, will remain fixed when possible. No chemistry will be violated as there are no additive modifications to the same site of a peptide sequence. In the event of additive effects being desirable, a corresponding Unimod entry can be added and employed (see also the utility add_unimod in Mzion).

### Supplementary Note 2

The considerable discrepancy in PSMs across search engines (Fig. 2c) prompted us to censor the search space of Mzion (psmC.txt). Taking the BI_G1 dataset as an example, nearly all PSMs from search engine Mascot have counterparts in psmC.txt whereas 5.4%, 0.8% and 0.5% of the PSMs from MaxQuant, MSFragger and MS-GF+, respectively, were not captured by Mzion (Table S1). Some of the extra findings can be ascribed to the side effects during the expansion of search space (Supplementary Note 1). Others can be explained by the insertion of complementary fragment ions^9^ to the original machine data during ion searches. For instance, with the approach, complementary b_2_, y_3_ and y_4_ and their doubly charged counterparts are assumed for a 7-residue peptide at machine data b_3_, b_4_ and y_5_. Mzion makes no assumption of ion complementarity. Moreover, it excludes the observations of secondary fragment ions *b*^0^, *y*^0^ etc. from being counted as independent evidence but uses merely their intensity information. PSMs with small numbers of primary ion matches will be excluded from psmC.txt at a default threshold of no less than six matched primary fragment ions.

We further examined the quality space of Mzion (psmQ.txt), illustrated with the BI_G1 dataset. The difference in the percentages of unidentified PSMs between psmC.txt and psmQ.txt suggests that approximately 1.4%, 7.4%, 6.2% and 2.9% of the PSMs that are absent from the psmQ.txt may be scoring-algorithm related when comparing Mzion to Mascot, MaxQuant, MSFragger and MS-GF+, respectively.

The absolute numbers in protein identifications are comparable across the search engines (Supplementary Fig. 3). In contrast, the differences in protein identities are relatively high (Table S1). In the example of dataset JHU_G1, 1.4% of peptides form Mascot were missing from psmQ.txt. However, the small percentage accounts for 15.5% of proteins that were lacking in Mzion. The imbalance between peptide and protein characterizations highlights the challenge in summarizing peptide evidences to protein identities, especially for proteins with low numbers of identifying peptides.^10^ One trait of Mzion is that it yielded generally fewer proteins with single-peptide identification (Supplementary Fig. 3b, Supplementary Fig. 4b and Supplementary Fig. 9b).

**Table S1**. Global PSMs, peptides, sequences and proteins from dataset WHIM_G. The definitions of peptide and sequence are in Fig. 1b.

|  | | **PSM** | |  | **Peptide** | |  | **Sequence** | |  | **Protein** | |
| --- | --- | --- | --- | --- | --- | --- | --- | --- | --- | --- | --- | --- |
| Set | Engine | ! Q% | ! C% |  | ! Q% | ! C% |  | ! Q% | ! C% |  | ! Q% | ! C% |
| BI_G1 | Mascot | 1.6 | 0.2 |  | 1.6 | 0.8 |  | 2.4 | 1.3 |  | 14.9 | 2.9 |
|  | MaxQuant | 12.8 | 5.4 |  | 6.0 | 2.0 |  | 5.8 | 2.1 |  | 19.2 | 4.0 |
|  | MSFragger | 7.0 | 0.8 |  | 6.7 | 1.7 |  | 8.8 | 3.3 |  | 11.1 | 3.6 |
|  | MS-GF+ | 3.4 | 0.5 |  | 3.8 | 1.4 |  | 4.7 | 2.1 |  | 13.9 | 4.8 |
| BI_G2 | Mascot | 1.5 | 0.2 |  | 1.5 | 0.9 |  | 2.3 | 1.4 |  | 14.2 | 2.9 |
|  | MaxQuant | 13.0 | 5.6 |  | 5.5 | 2.0 |  | 5.4 | 2.1 |  | 17.9 | 4.0 |
|  | MSFragger | 6.3 | 0.7 |  | 6.1 | 1.7 |  | 8.4 | 3.3 |  | 10.7 | 3.8 |
|  | MS-GF+ | 3.1 | 0.5 |  | 3.5 | 1.5 |  | 4.3 | 2.1 |  | 12.5 | 4.7 |
| JHU_G1 | Mascot | 1.3 | 0.2 |  | 1.4 | 0.9 |  | 2.3 | 1.5 |  | 15.5 | 3.6 |
|  | MaxQuant | 12.0 | 5.7 |  | 4.4 | 1.8 |  | 4.2 | 1.9 |  | 18.6 | 4.5 |
|  | MSFragger | 5.8 | 0.7 |  | 5.4 | 1.6 |  | 8.1 | 3.7 |  | 11.1 | 4.3 |
|  | MS-GF+ | 2.7 | 0.4 |  | 3.0 | 1.4 |  | 4.0 | 2.2 |  | 13.1 | 5.5 |
| JHU_G2 | Mascot | 1.6 | 0.2 |  | 1.5 | 0.8 |  | 2.5 | 1.5 |  | 16.0 | 3.3 |
|  | MaxQuant | 16.2 | 7.3 |  | 6.1 | 2.2 |  | 5.8 | 2.3 |  | 19.6 | 4.4 |
|  | MSFragger | 6.2 | 0.7 |  | 5.6 | 1.5 |  | 8.5 | 3.7 |  | 12.2 | 4.1 |
|  | MS-GF+ | 2.8 | 0.4 |  | 3.0 | 1.3 |  | 4.1 | 2.0 |  | 14.0 | 5.3 |
| PNNL_G1 | Mascot | 2.2 | 0.3 |  | 2.1 | 1.2 |  | 3.2 | 1.8 |  | 15.2 | 3.4 |
|  | MaxQuant | 15.6 | 7.4 |  | 5.4 | 2.0 |  | 5.3 | 2.1 |  | 18.9 | 4.5 |
|  | MSFragger | 7.9 | 0.8 |  | 7.0 | 2.1 |  | 9.9 | 4.1 |  | 12.7 | 4.3 |
|  | MS-GF+ | 3.9 | 0.5 |  | 4.2 | 1.7 |  | 5.5 | 2.5 |  | 13.7 | 5.3 |
| PNNL_G2 | Mascot | 2.3 | 0.3 |  | 2.3 | 1.2 |  | 3.4 | 1.9 |  | 15.4 | 3.4 |
|  | MaxQuant | 15.8 | 7.5 |  | 5.7 | 2.1 |  | 5.5 | 2.2 |  | 19.0 | 4.3 |
|  | MSFragger | 8.0 | 0.8 |  | 7.1 | 2.1 |  | 10.1 | 4.1 |  | 12.6 | 4.1 |
|  | MS-GF+ | 3.9 | 0.5 |  | 4.3 | 1.7 |  | 5.6 | 2.5 |  | 13.5 | 5.2 |
| ^1^ ! Q% - Percent not in the quality space of psmQ.txt | | | | | | | | | | | | |
| ^2^ ! C% - Percent not in the complete space of psmC.txt | | | | | | | | | | | | |

### Supplementary Note 3

#### The ion statistics with Mzion.

In the previous note, we inspected PSMs and peptides that were missing from the Mzion space (Supplementary Note 2). To further understand the behaviors of Mzion, we characterize additionally the ion statistics of PSMs using the six datasets in WHIM_G.

We first investigated the relative average of MS1 intensity that are unique to each search engine, namely, $\bar{I_{ui}}/\bar{I_{ai}}$ where $I_{ui}$ is the unique intensity and $I_{ai}$ the overall intensity by a search engine $i$. Here overall intensity refers to the summed MS1 signals from all significant PSMs (sigPSM) by a search engine and unique intensity refers to the summed MS1 signals from the sigPSM subsets that were uniquely determined by a search engine (Fig. 2c). The absolute MS1 intensity varied by search engines partially due to the peak representation, for example, by either heights or areas; thus the relative intensity was used. Among the six datasets, we found that Mzion gave the highest relative mean intensity in unique sigPSM (Fig. S1a). Analogous analysis against MS2 reporter ions showed that Mzion also yielded the greatest relative mean in unique reporter-ion intensity (Fig. S1b). Additional difference in the sigPSM space includes the populations of precursor *m*/*z* values where Mzion tends to capture more features concentrated around *m*/*z* 600 and 700 (Fig. S1c). The range is typically associated with high detection sensitivities in ion-trap MS, which might explain the high relative means in both precursor and reporter-ion intensities with Mzion.

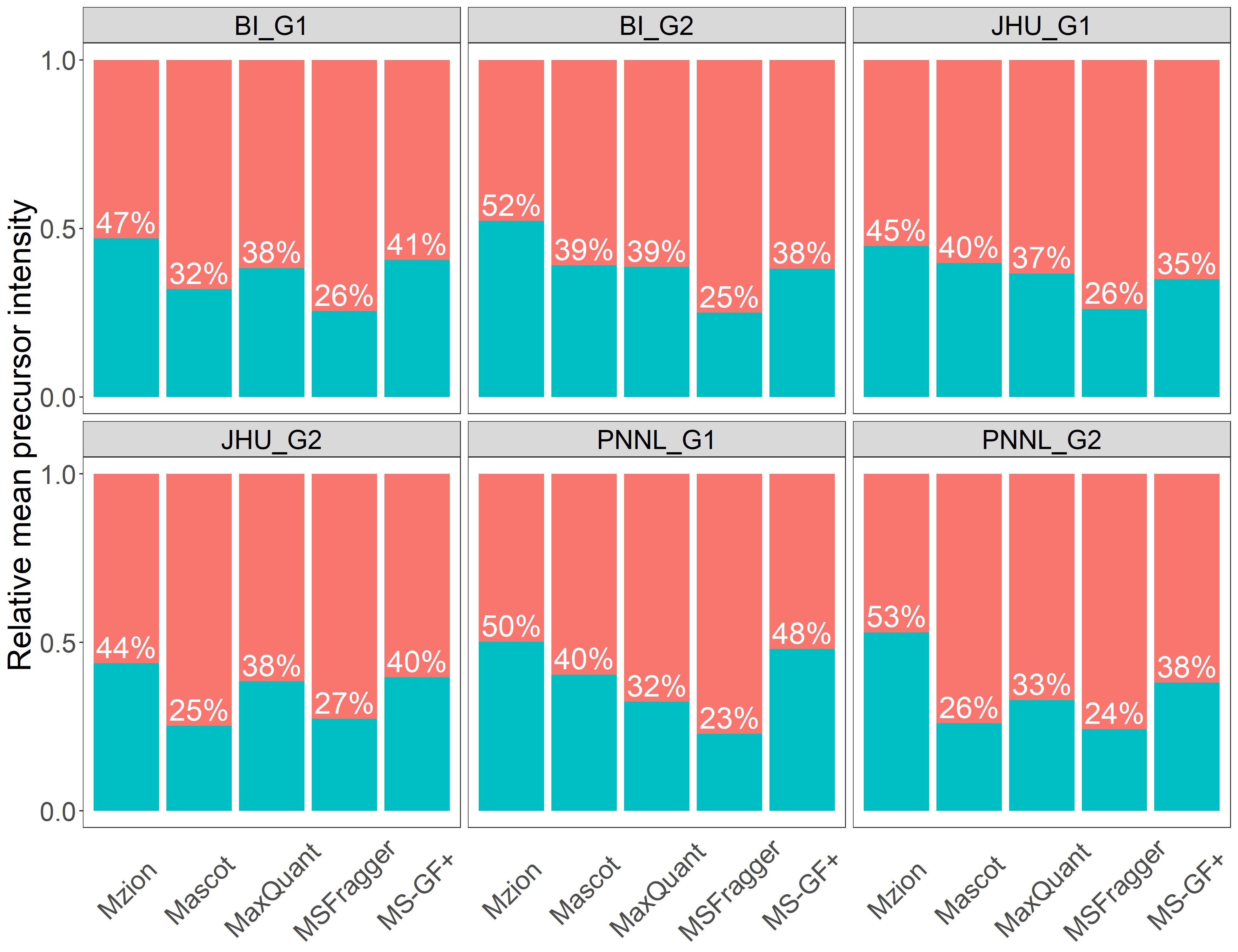
**Figure S1.** (a) The proportion of engine-specific (unique) precursor intensity from dataset WHIM_G. (b) Mean statistics of unique reporter-ion intensity from dataset WHIM_G. (c) Distributions of unique precursor *m*/*z* from dataset WHIM_G.

**a**

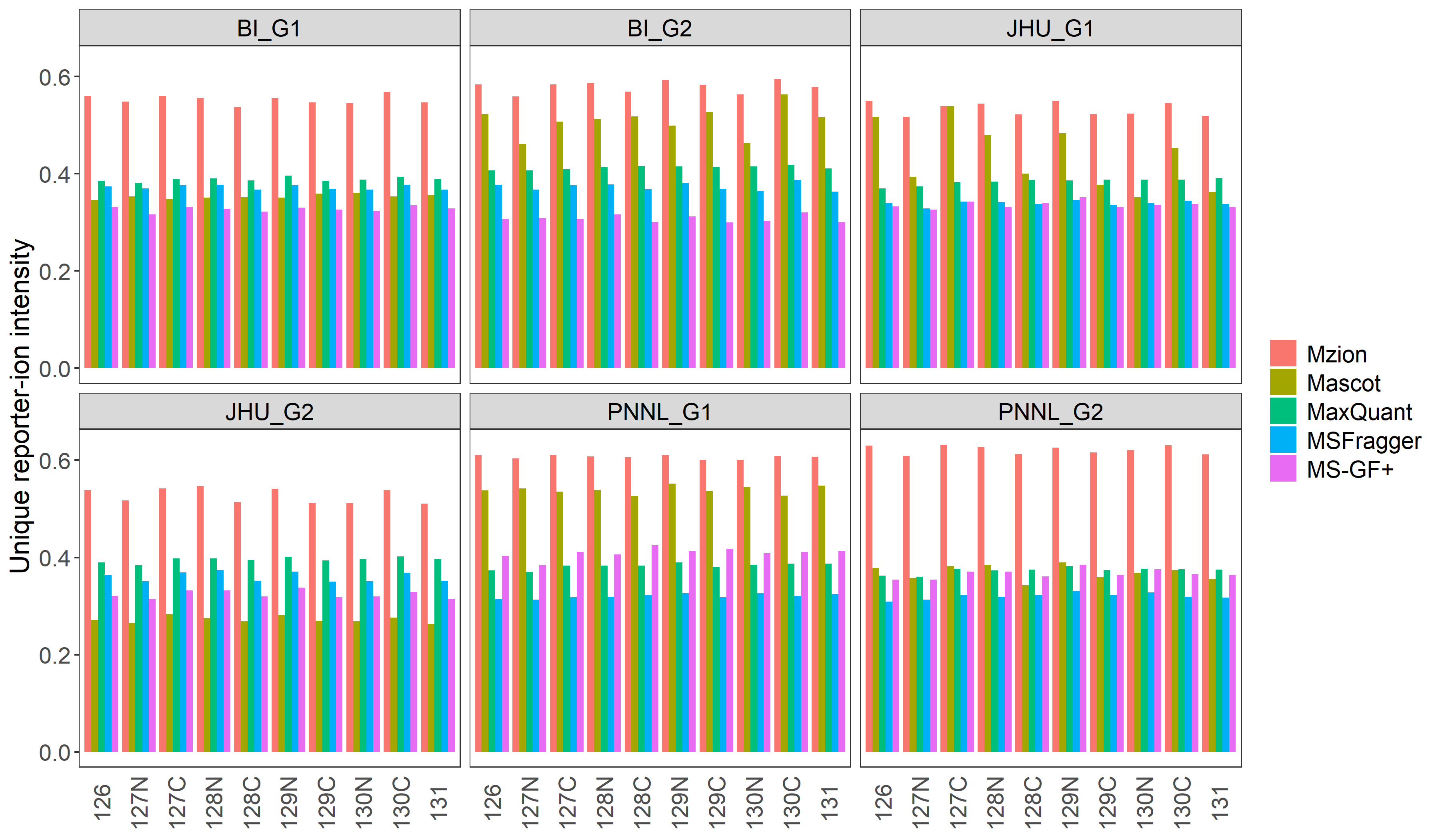

**c**

**b**

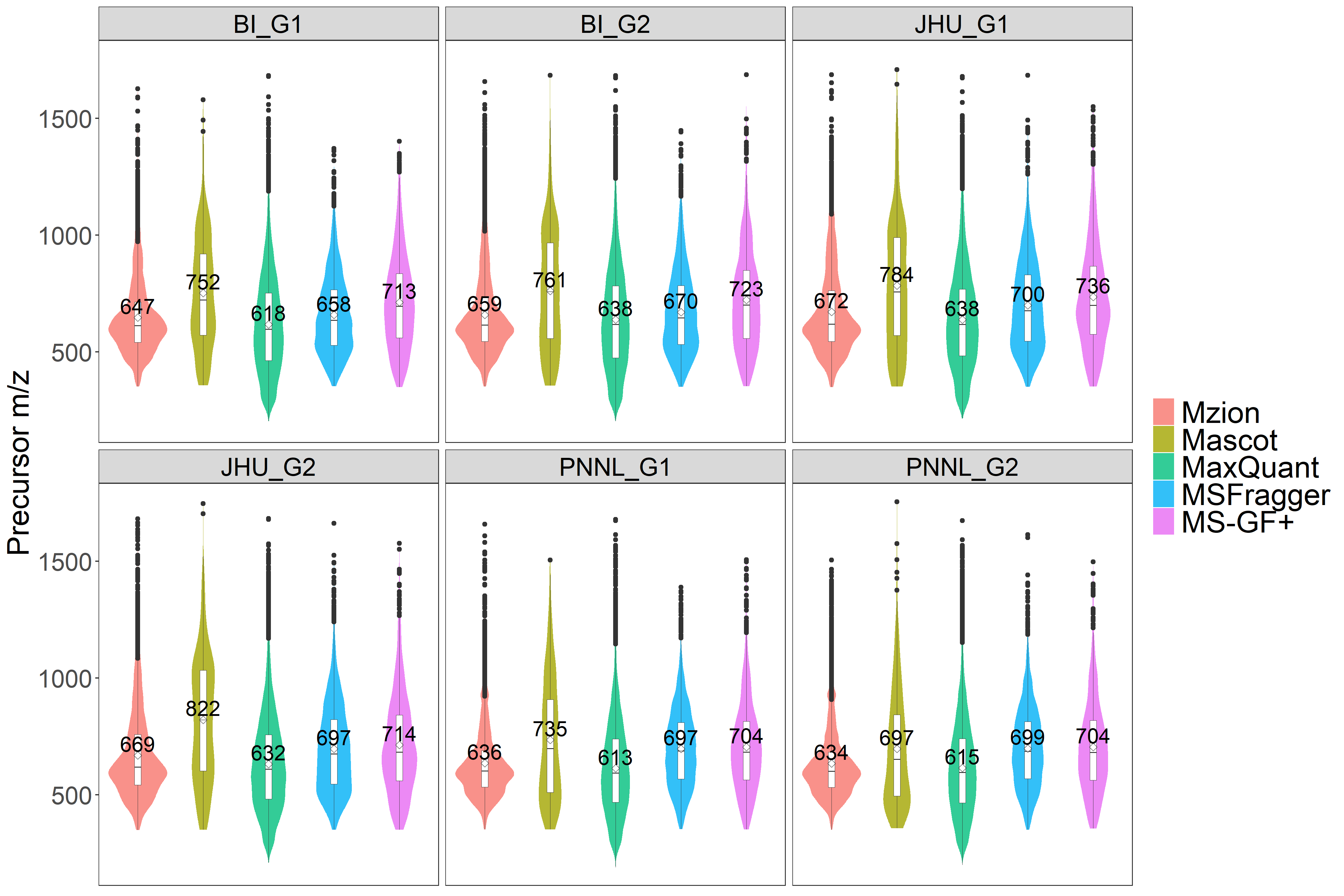

### Supplementary Note 4

#### Correlations and distances between replicated samples

One caveat of Pearson’s correlation ($\rho$) is its sensitivity to extreme values. The behavior can be illustrated with a simulated example. Note that mass spectroscopic readouts are recorded as intensity values. To convert intensity to the relative intensity of log2FC in conventional proteomic procedures, it requires references for uses as denominators. We first generated three vectors of random intensity values, $I_{x}$, $I_{y}$ and $I_{r}$ ($\in\mathbb{R}^{1000}$) ranging from 1E3 to 1E7, where $I_{y}$ and $I_{r}$ are correlated to $I_{x}$ by normal errors. We next converted $I_{x}$ and $I_{y}$ to log2FC, $\mathbf{x}$ and $\mathbf{y}$, with $I_{r}$ being the reference, namely, $\mathbf{x}=\text{log}_{2}\left( I_{x}/I_{r} \right)$ and $\mathbf{y}=\text{log}_{2}\left( I_{y}/I_{r} \right)$. With the setup, $\mathbf{x}$ and $\mathbf{y}$ were correlated with $\rho=0.715$. If we were to exclude the five greatest and the five least values from $\mathbf{x}$, the $\rho$ drops noticeably to $0.701$ (Fig. S2a).

The above example alluded to an effect of reference choices on the correlation of data. For instance, in the event of a vast difference between samples and references, extreme values at high or low log2FC can be introduced during the intensity-to-log2FC transformation. As a result, elevated corrections can be expected. To illustrate, we took randomly 20 values from the reference and multiplied them by 0.2. With the new $I_{r}$ but the same $I_{x}$ and $I_{y}$, the correlation between $\mathbf{x}$ and $\mathbf{y}$ are now enhanced substantially to $0.926$ (Fig. S2b).

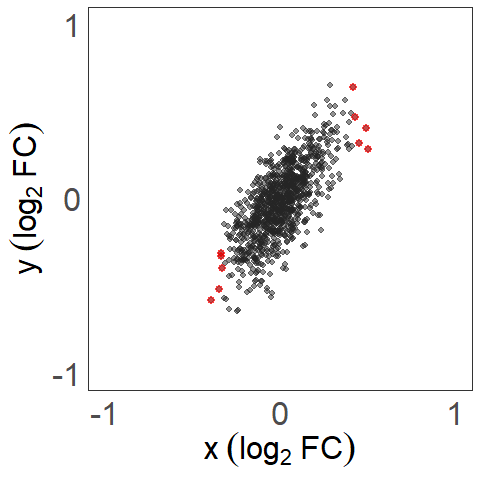

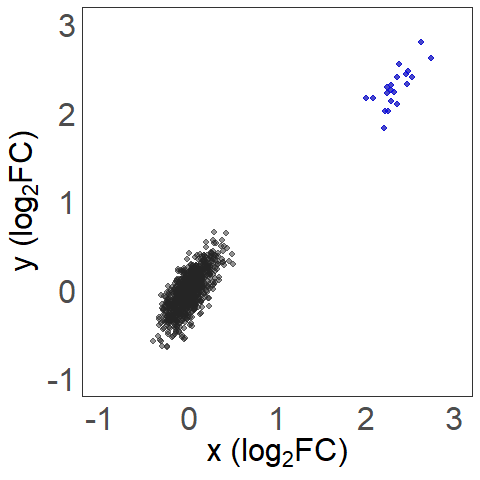
**Figure S2**. Simulated correlations. (a) Removal of extreme values. (b) Introduction of extreme values.

**b**

**a**

In the above examples, we correlated $I_{y}$ and $I_{r}$ to $I_{x}$ with independent random errors. We next simply exchange the values of $I_{x}$ and $I_{r}$. As a result, $I_{x}$ and $I_{y}$ are each correlated *independently* to $I_{r}$. The corresponding log2FC, $\mathbf{x}$ and $\mathbf{y}$, are thus independently and identically distributed (iid) variables. As expected, the newly calculated $\rho=-0.035$ is close to the theoretical expectation of 0 (Fig. S3a). The scenario of high variability in correlations can be tied to the examples of WHIM2 (basal) and WHIM16 (luminal) replicates in the BI_G1 dataset. That is, a high correlation was observed when comparing two WHIM2 replicates with respect to the average of all WHIM2 and WHIM16 (Fig. 2d) whereas a near zero correlation was found for the same pair of WHIM2 with respect to the average of all five WHIM2s (Fig. S3b).

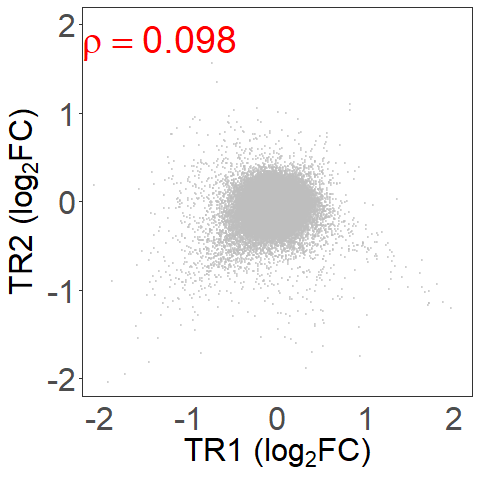

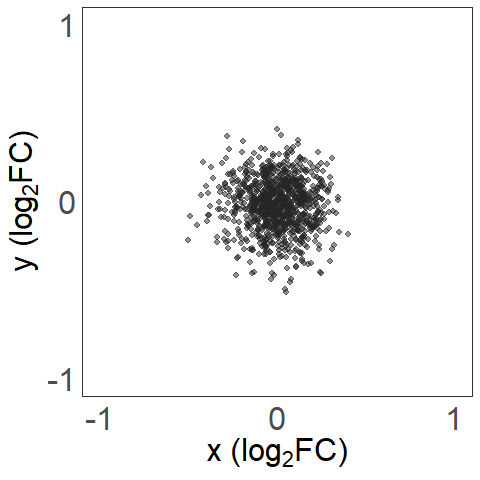
**Figure S3**. Correlations of iid variables. (a) Simulated. (b) The first two WHIM2 replicates from dataset BI_G1.

**b**

**a**

For the above reasons of high sensitivity to reference choices, we assessed additionally the data quality by the Manhattan distance, $\delta=\frac{1}{n}\sum_{i=1}^{n} \left( \left| x_{i}-y_{i} \right| \right)$ where $x$ and $y$ are log2FC and $n$ is the number of data pairs. A shorter distance indicates a greater similarity between samples. Different to $\rho$, the distance depends less on the choices of references, albeit a missing value in reference will nullify the corresponding {$x_{i},y_{i}$} pair and decrease $n$ by one.

#### Choice of samples and references

We focus arbitrarily on the comparison of WHIM2 instead of WHIM16 when assessing pairwisely the Manhattan distance between replicates. We noted that the use of reference WHIM2 alone (5 samples) often yielded fewer missing values than the use of combined reference WHIM2+WHIM16 (10 samples). The larger extent of missing values with the latter can be attributed to the considerable difference between the two breast cancer subtypes. In other words of $n$ pairs of data {$x_{i},y_{i}$} ($i=1,2,...,n$), a greater disparity between $x_{i}$ and $y_{i}$ increases the chance of a missing value in either $x_{i}$ or $y_{i}$, which subsequently trivializes the contribution from the {$x_{i},y_{i}$} pair and decreases $n$ by one. To utilize more of the available information, the reference WHIM2 was chosen in distance assessments.

Among the six datasets in WHIM_G, the first two WHIM2 replicates often yielded shorter distances (D12) than other pairs (Supplementary Fig. 5b). Presumably, a shorter distance is associated with smaller technical artefacts. To focus on the aspect of data similarity that can be conferred by search engines, we chose the first two replicates, TR1 and TR2, for the demonstration of global data (Fig. 2d, Supplementary Fig. 5a). By the same token, we enumerated primarily replicates TR1 and TR2 from dataset WHIM_P when demonstrating phosphopeptide data (Fig. 3, Supplementary Fig. 7).

### Supplementary Note 5

#### Entrapment by database entries

The analyses were performed against the six global datasets in WHIM_G at 1% protein FDR, with the exception of Mascot and MS-GF+ results at an engine-defined 1% PSM FDR. The target database includes RefSeq proteins at species human mouse (Jul. 2013; 56,673 entries) and common contaminants (cRAP, Jan. 2012; 116 entries). The entrapment database is Uniprot arabidopsis thaliana (Sept. 2022; 16,312 entries). PSMs were sorted by decreasing probability scores. Peptides or sequences were described with their best-scored PSM. Peptides that can be found from both target and entrapment databases were flagged as targets when calculating the entrapment rates. Proteins were sorted by the descending numbers of unique identifying sequences. The entrapment rates were reported as the ratio of the number of entrapped identities divided by the total number of identities at each of the PSM, peptide, sequence and protein levels.

In general, all search engines performed comparably in the above four categories (Fig. S4). Surges in entrapment around the onset of the rank plots were typically observed. The early rises can be ascribed to the assignments of tandem spectra to entrapment peptides with high probability scores. In the example of dataset BI_G1, the highest-score entrapment peptide by Mzion is AAAASLLGK under protein GRV2 at accession F4IVL6|GRV2_ARATH. For an unknown reason, MSFragger (version 18.0) assigned additionally the peptide to target protein DDTL at accessions NP_001346 that lacks the sequence. To have a more common estimate of entrapment rates across search engines, we forced peptides that were identified as decoys by any search engine to decoy entries.

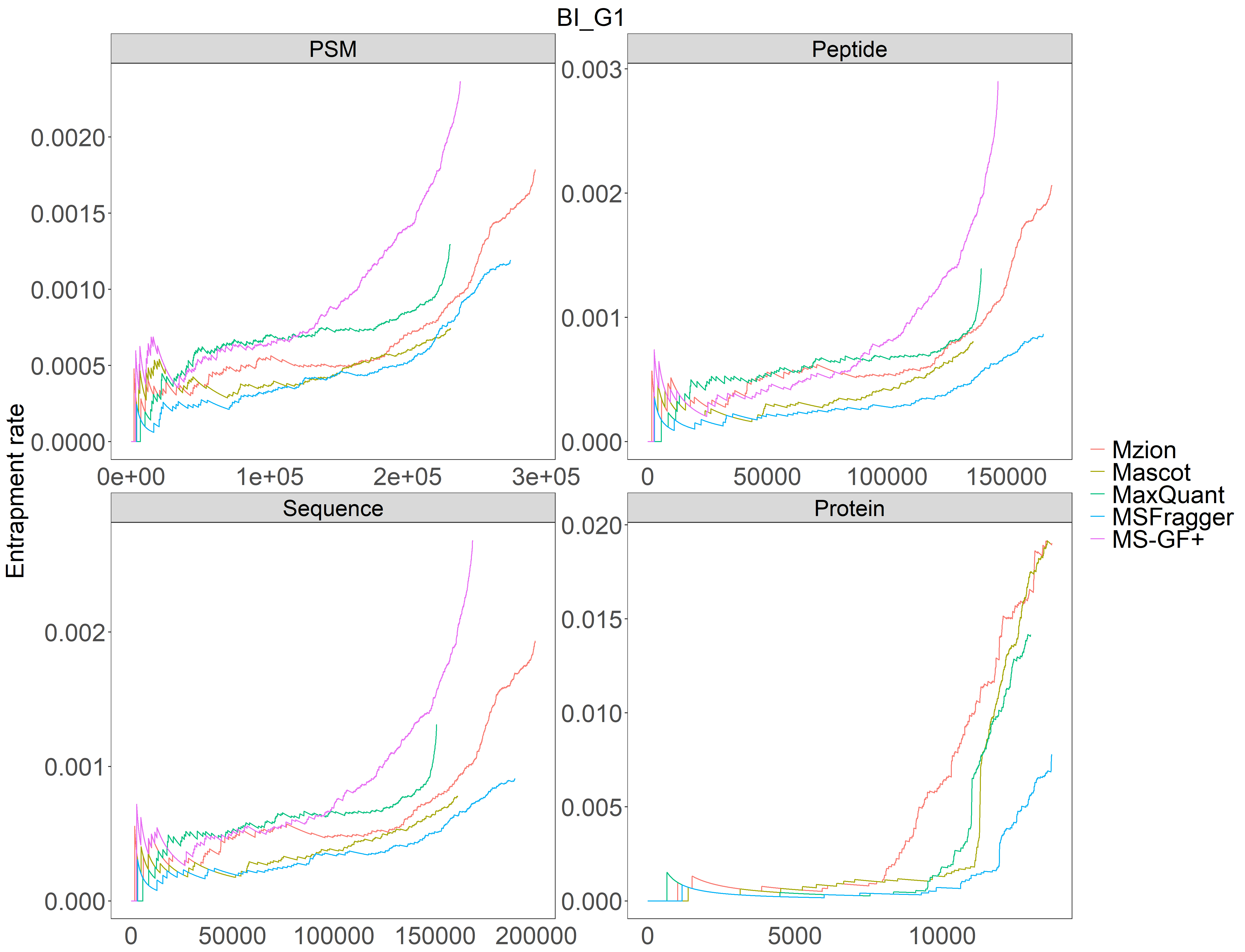
**Fig. S4**. Entrapment by database (arabidopsis thaliana) entries from dataset WHIM_G.

**a**

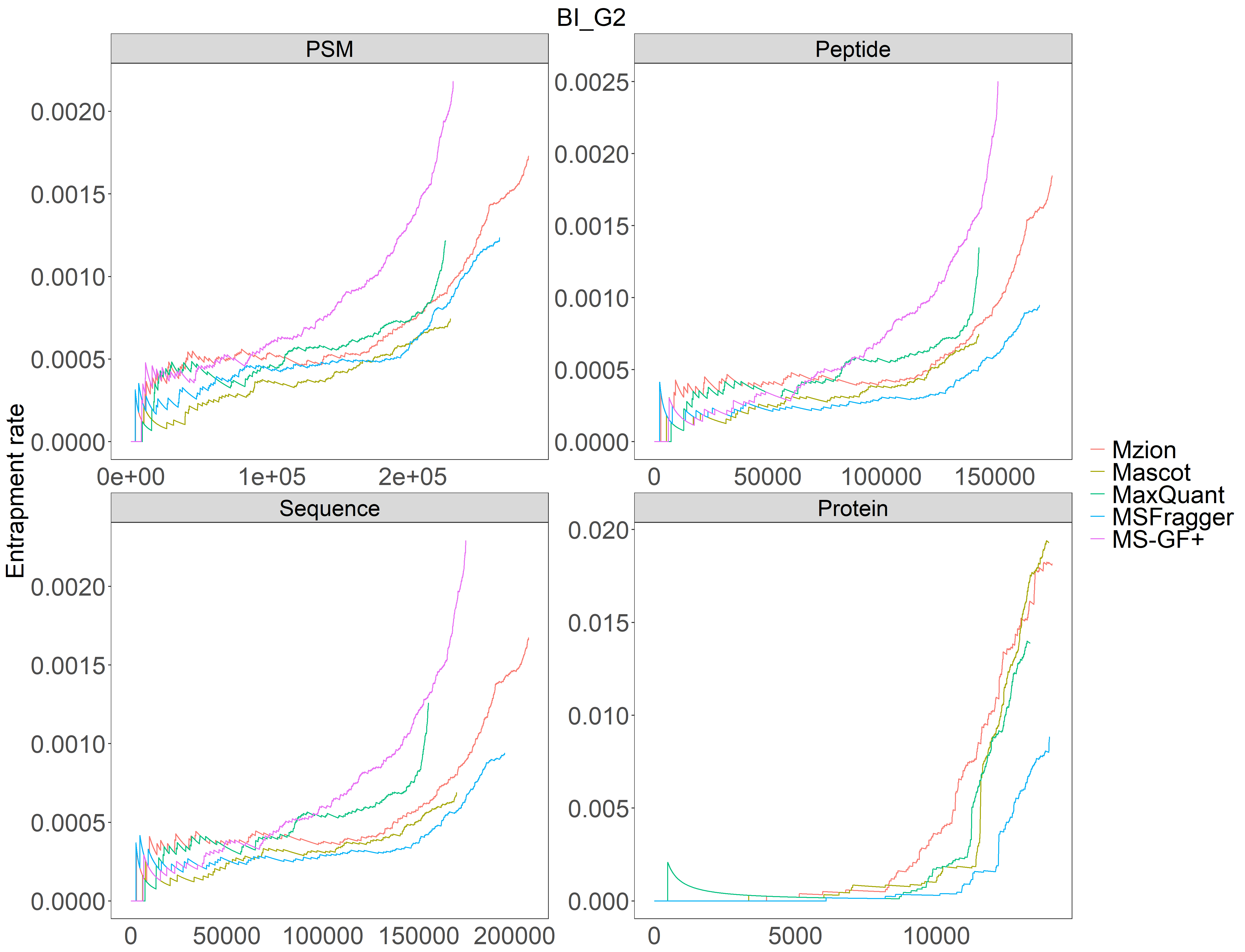

**b**

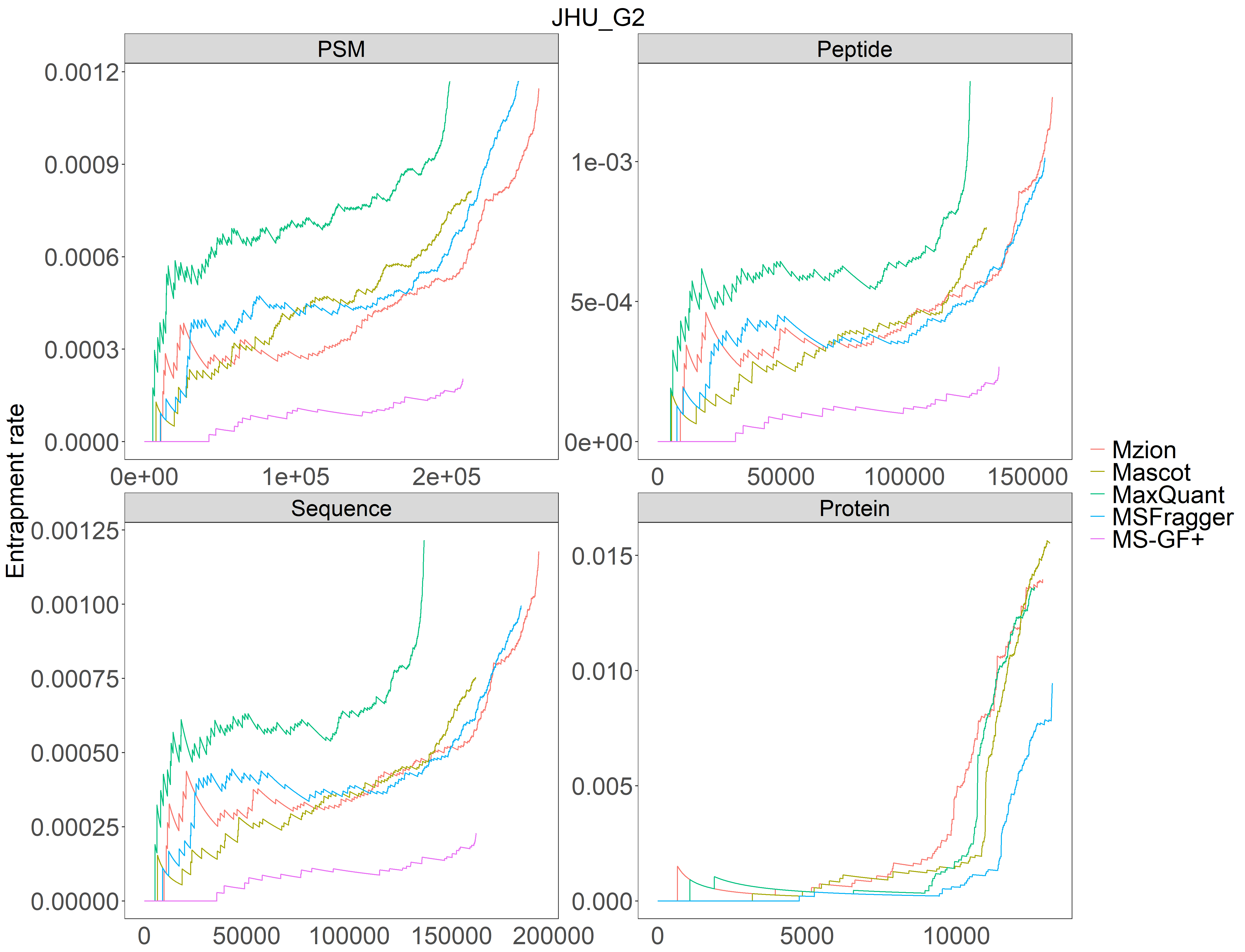

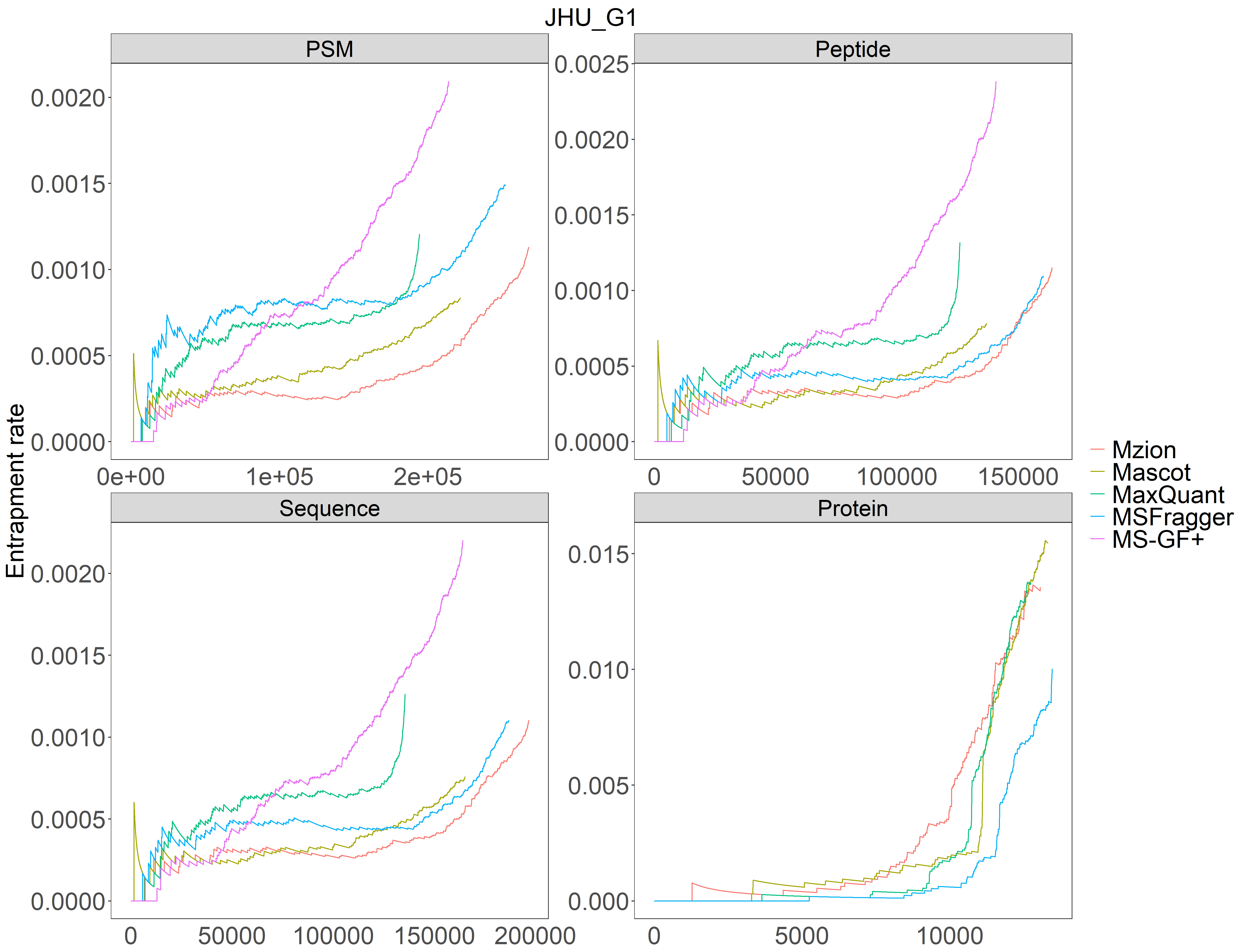

**d**

**c**

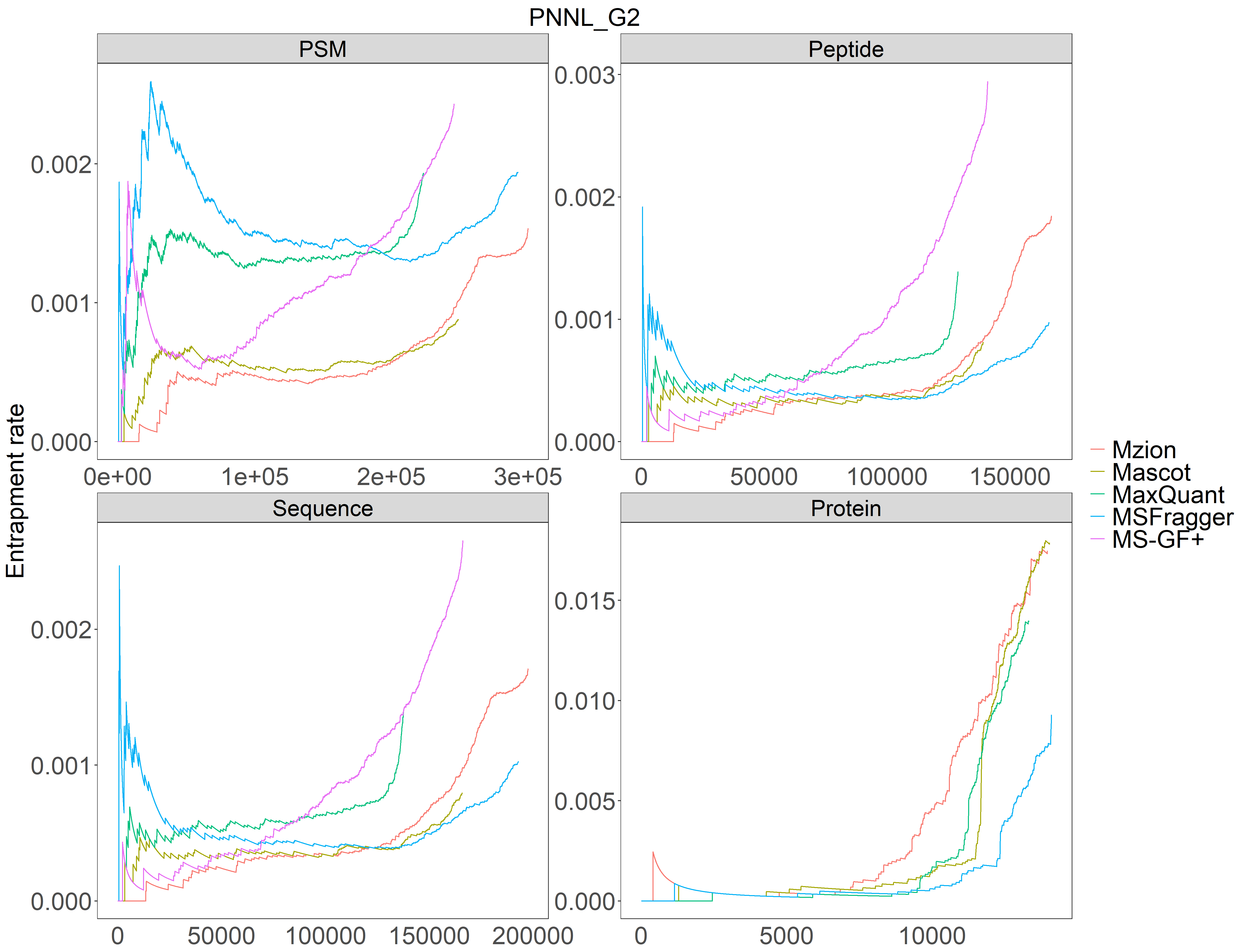

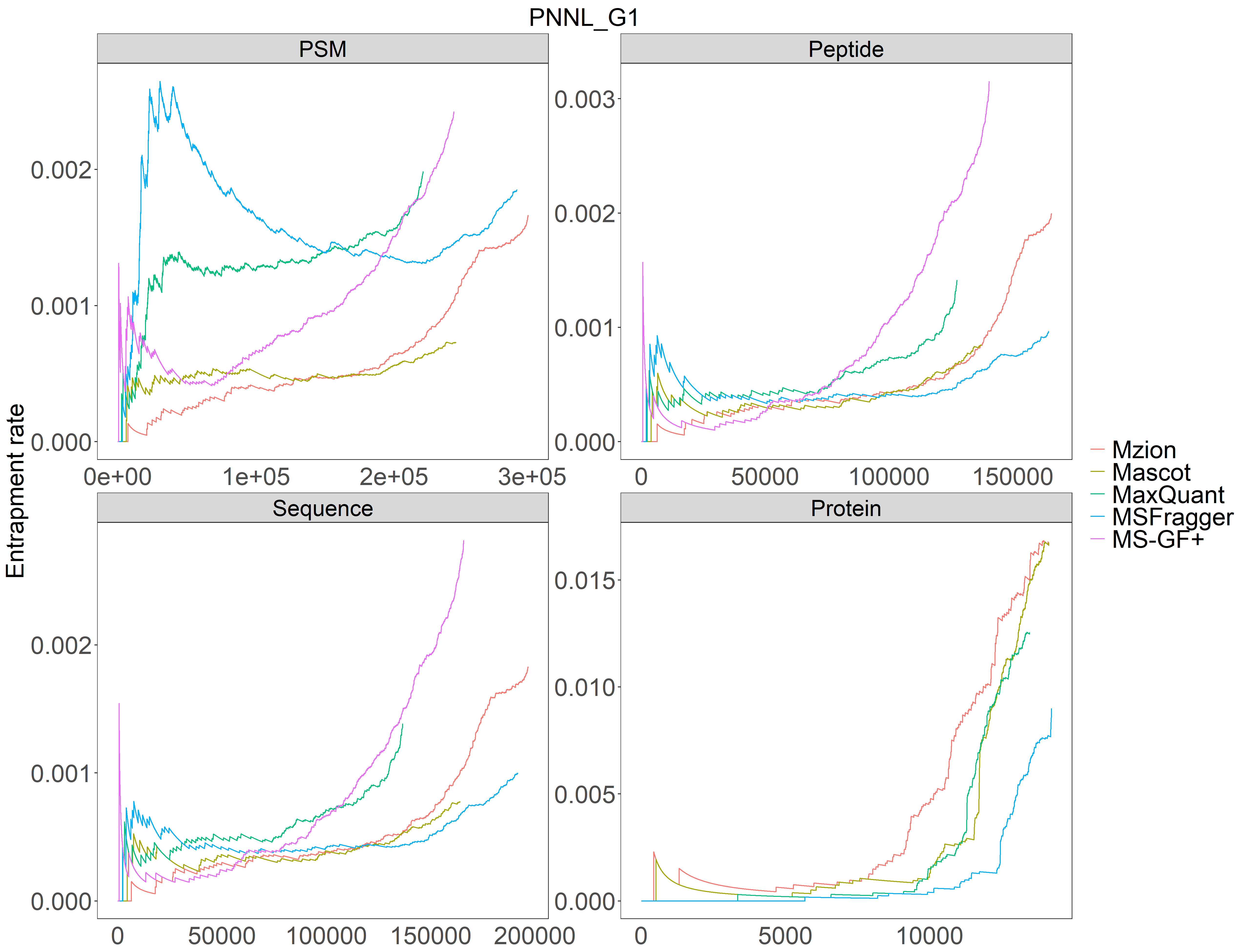

**e**

**f**
